# Modeling reveals cortical dynein-dependent fluctuations in bipolar spindle length

**DOI:** 10.1101/2020.07.10.197285

**Authors:** Dayna L. Mercadante, Amity L. Manning, Sarah D. Olson

## Abstract

Proper formation and maintenance of the mitotic spindle is required for faithful cell division. While much work has been done to understand the roles of the key molecular components of the mitotic spindle, identifying the consequences of force perturbations in the spindle remains a challenge. We develop a computational framework accounting for the minimal force requirements of mitotic progression. To reflect early spindle formation, we model microtubule dynamics and interactions with major force-generating motors, excluding chromosome interactions that dominate later in mitosis. We directly integrate our experimental data to define and validate the model. We then use simulations to analyze individual force components over time and their relationship to spindle dynamics, making it distinct from previously published models. We show through both model predictions and biological manipulation that rather than achieving and maintaining a constant bipolar spindle length, fluctuations in pole to pole distance occur that coincide with microtubule binding and force generation by cortical dynein. Our model further predicts that high dynein activity is required for spindle bipolarity when kinesin-14 (HSET) activity is also high. Together, our results provide novel insight into the role of cortical dynein in the regulation of spindle bipolarity.

**SIGNIFICANCE:** The mitotic spindle is a biophysical machine that is required for cell division. Here we have paired a modeling approach with experimental data to understand the maintenance and dynamics of a bipolar mitotic spindle in the absence of chromosome interactions. We present novel roles of cortical dynein in mitosis, and demonstrate its requirement for both dynamic changes in spindle length and in antagonizing HSET in bipolar spindle formation. Model outputs predict that cortical dynein activity would be limiting in contexts where HSET activity is high and may be of therapeutic relevance in cancer contexts where HSET is often over expressed.

## INTRODUCTION

Mathematical and computational modeling of biological processes can bypass experimental limitations and provide a framework to identify and manipulate individual molecular components. An appealing candidate for such modeling is the process of cell division (1), which involves formation of the mitotic spindle to organize and separate the genetic material of a cell into two identical daughter cells. The assembly of the mitotic spindle is initiated by the nucleation of microtubules (MTs) at an organelle known as the centrosome (2). Normal mitotic cells have two centrosomes at which the spindle poles are formed. The centrosomes are positioned in response to mechanical forces, primarily driven by the activity of motor proteins (3–7). As mitosis proceeds, the mitotic spindle forms and maintains a bipolar configuration, with the two centrosomes positioned at opposite sides of the cell.

Many models have been developed to understand early centrosome separation and spindle formation (8–15), chromosome dynamics (16–21), and spindle elongation during anaphase (22–24). While varying widely in methods and biological motivation, computational force-balance models have been used to understand key mechanistic components that modulate positioning of spindle poles and bipolar spindle formation (13, 25–28). Due to the ambiguity surrounding the exact spatiotemporal distribution and motor force generation in cells, and the large number of MT-motor interactions (6, 29, 30), computational models generally simplify dynamics and focus on the role of a limited number of interactions. We also use a simplified approach to modeling motor-MT interactions, where, rather than modeling each individual motor protein in time and space, we set a probability that a motor protein will stochastically bind and generate force based on its proximity to a MT or the cell boundary (13, 27).

Proper formation of the mitotic spindle is required for accurate chromosome segregation, and while the molecular regulation of segregation onset is dependent on stable MT attachments to chromosomes (31), chromosomes are dispensable for early bipolar spindle assembly (32–34). Hence, we develop a minimal computational model to analyze centrosome movement and mammalian mitotic spindle formation in the absence of chromosomes. To better understand the key mechanistic requirements of bipolar spindle formation and maintenance in the absence of stable MT interactions with chromosomes, we explore how forces drive centrosome movement. We consider stochastic MT interactions and forces generated by three motor proteins: kinesin-5 (Eg5), kinesin-14 (HSET), and dynein, which have been extensively studied and identified as the major force generators in mitosis (7, 35, 36). We leverage prior molecular studies that have identified velocity, force, and force scales for motor proteins to define parameters for our model (37–41).

Discerning the distinct role(s) of motor-dependent forces on mitotic progression has been challenging as some mitotic motors have two or more regions of localization and/or functions that are independently regulated in the cell (7, 36). Dynein, for example, is localized to and interacts with MTs at spindle poles, kinetochores, and the cell cortex (36). Cell biological approaches can be limited in their ability to selectively perturb one localization or function of this important motor. Here we model cortical- and spindle pole-localized dynein independently, allowing us to assess the force generation of each population separately. Our model also explores temporal changes in motor-dependent forces and their impact on spindle dynamics. By analyzing motor-dependent force generation through mitotic progression, we answer outstanding questions regarding the balance of forces during cell division. Specifically, we test the impact of cortical dynein activity on spindle bipolarity and explore how force perturbations impact bipolar spindle length in the absence of cortical dynein. We directly integrate fixed and live-cell imaging to both refine and validate model outputs. Our model captures the biological time scale of mitotic progression and recapitulates changes to bipolar spindle length that have been previously described following molecular or genetic perturbation of motor protein function. Using our model, together with cell biological analysis of dynein perturbation, we reveal that cortex-localized dynein impacts both bipolar spindle length and spindle length fluctuations during mitosis. Model results further indicate that cortical dynein activity antagonizes HSET-derived forces on antiparallel MTs to directly impact bipolar spindle length.

## MATERIALS AND METHODS

### Model Overview

In our two-dimensional simulations, the cell cortex is a rigid, circular boundary, with a diameter of 30 *µ*m, capturing a mammalian cell that has rounded as it enters mitosis (42–44). We allow MT-motor protein interactions with Eg5 and HSET on antiparallel MTs, capturing the dominant roles of these proteins in mitosis, and with dynein at the cell cortex and spindle poles (4, 6, 7, 25, 30, 35, 45, 46). We use a simplified approach to determine MT interactions and force generation based on a Monte Carlo binding probability. Hence, for computational simplicity, we do not model individual motor proteins and do not include chromosomes, kinetochores or kinetochore fibers, as these are dispensable for bipolar spindle formation and maintenance (Fig. 3). Where available, experimentally defined parameters using mammalian cell culture were used, and all parameters described below are listed in Table S1 in the Supporting Material. The model is benchmarked on previous modeling approaches that capture dynamic centrosome positioning and cell division (13, 27, 47–49). Additional model validation and details are provided in the Supporting Material.

**Figure 1:**
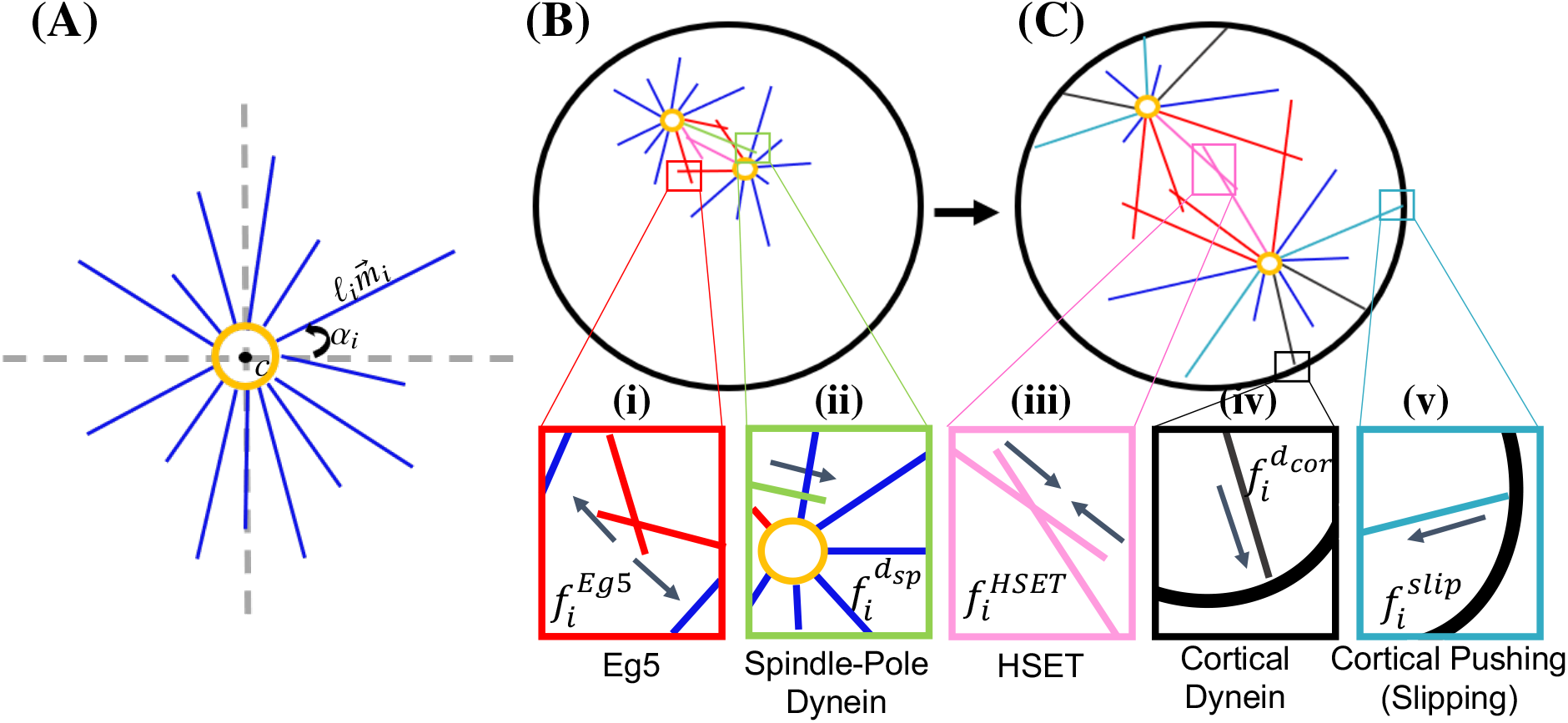
MT-associated forces are involved in forming and maintaining a bipolar spindle in the absence of chromosome interactions. (A) MTs are nucleated from and remain anchored at centrosome *c*. Each MT *i* is defined by an angle *α*_*i*_, length *ℓ*_*i*_ and direction 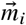. A balance of pushing and pulling forces are required for proper centrosome separation in (B) and maintenance of spindle bipolarity in (C). (i) Eg5 generates an outward force at antiparallel MT overlap regions. (ii) Dynein localized to spindle poles binds to and pulls MTs from the opposing spindle pole. (iii) HSET generates an inward force at antiparallel MT overlap regions. (iv) Dynein localized to the cell cortex generates a pulling force on bound MTs. (v) MTs that continue to grow as they reach the cell cortex generate a pushing force. Arrows in (i)-(v) indicate the direction in which centrosome *c* will move in response to *f*_*i*_.

**Figure 2:**
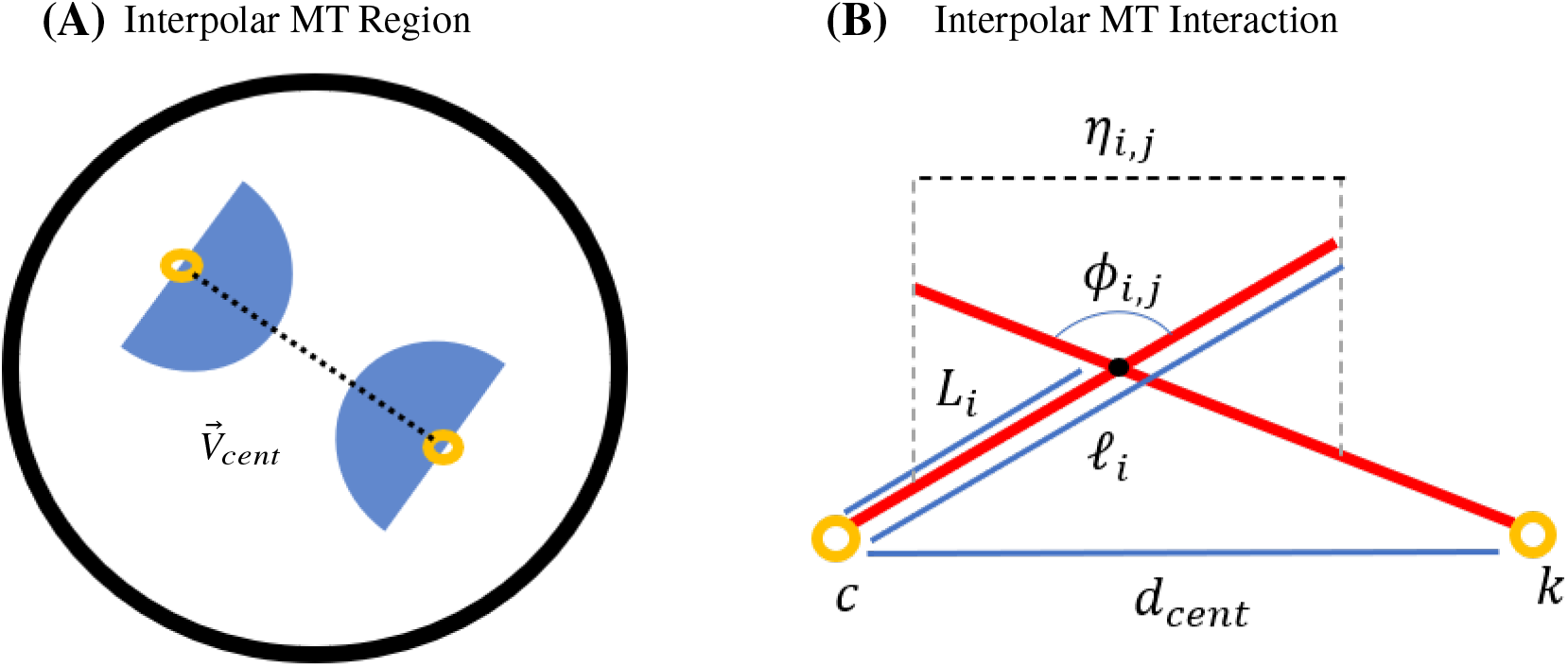
Interpolar MTs. (A) Schematic of interpolar MT region. Black dashed line indicates the vector between the centrosomes 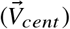. Interpolar MTs are those that lie within the blue shaded regions. (B) Schematic of interpolar MT interaction. MTs *i, j* are nucleated from centrosomes *c, k*, respectively.

**Figure 3:**
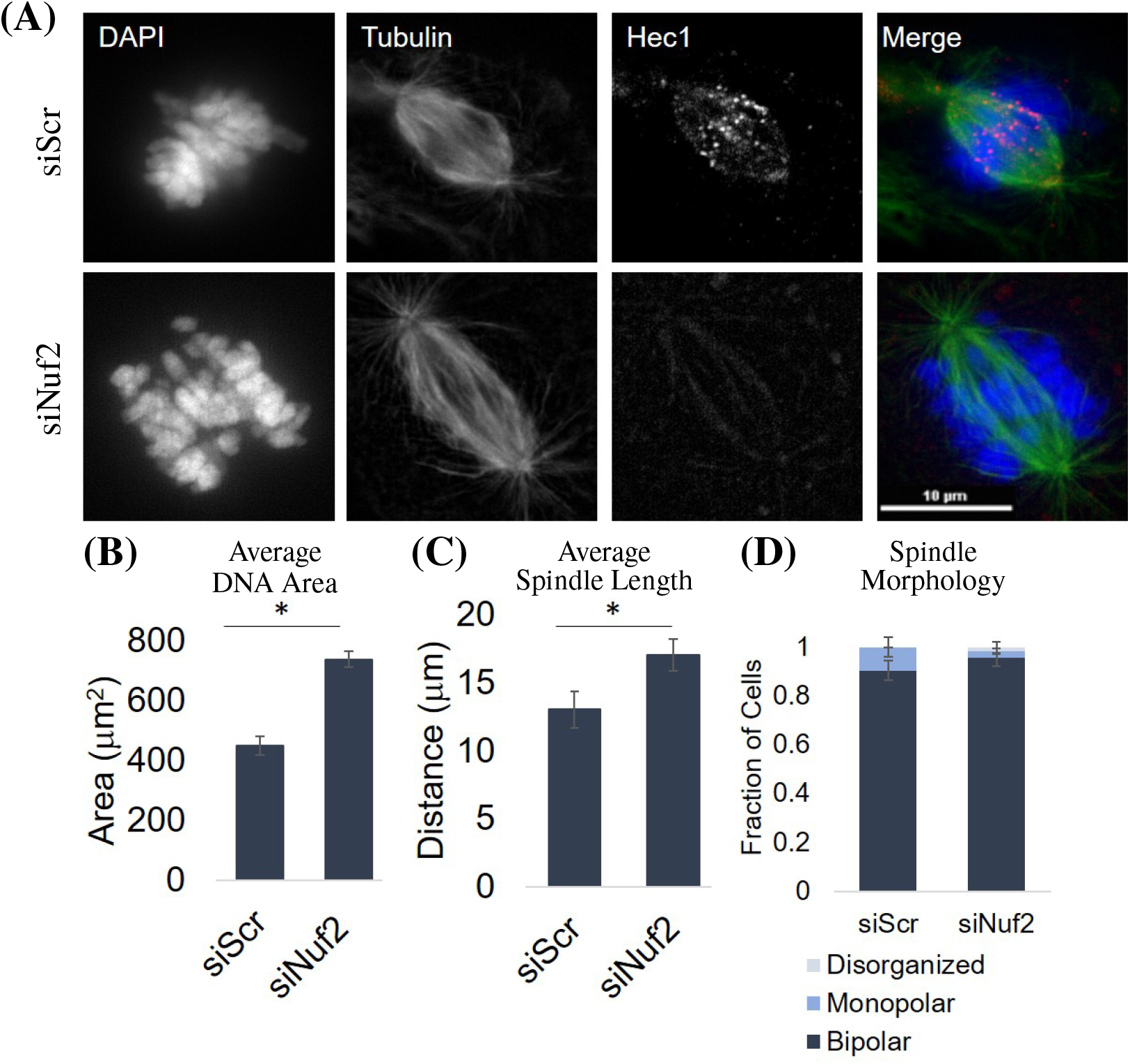
Stable end-on kinetochore attachments are not required for bipolar spindle formation. (A) Fixed-cell imaging of RPE cells stained for DAPI (DNA), Tubulin (MTs), and Hec1 (Ndc80 complex) in the control (siScr) and knockdown (siNuf2) condition. (B) Quantification of the average DAPI area in the control (siScr) and knockdown (siNuf2) condition. (C) Quantification of the average spindle length in the control (siScr) and knockdown (siNuf2) condition. (D) Quantification of the average fraction of pre-anaphase mitotic cells cells with bipolar, monopolar, or disorganized spindles. All averages were calculated from at least 30 cells from 3 biological replicates. Error bars are standard deviation (SD). *p*<*0.05 indicates statistical significance.

### Dynamic Microtubules

MTs are elastic filaments oriented such that their plus-ends, those that dynamically grow and shrink (50), point outward while their minus ends remain anchored at the centrosome (51–53). We consider MT minus-ends to remain embedded in the centrosome (*c* in Fig. 1 A) to account for crosslinking proteins that maintain spindle-pole focusing throughout mitosis (54, 55). MT plus-ends undergo dynamic instability (50), meaning that they are stochastically switching between states of growing (at a velocity *v*_*g*_) and shrinking (at a velocity *v*_*s*_ if unbound or *v*_*b*_ if bound to cortical dynein). Each MT *i* is nucleated from one of the two centrosomes, has an angle *α*_*i*_, length *ℓ*_*i*_, and is characterized by a unit direction vector 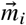 from the center of the centrosome to the MT plus-end (Fig. 1 A). As MTs interact with each other or the cell boundary, 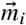 further defines the direction in which motor-dependent forces are generated on MT *i*, and therefore felt on centrosome *c*.

### Centrosome Movement

We define five MT-derived forces that drive the movement of centrosome *c* within the confined cell boundary; pushing forces by MTs growing against the cell cortex 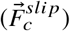, motor-dependent pulling forces by dynein at the cell cortex 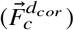 or spindle poles 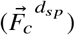, and Eg5-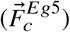 or HSET-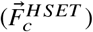 derived forces at interpolar MT overlap regions (Fig. 1 B/C (i)-(v)). Since exact amounts and distributions of motor proteins throughout the spindle have not been experimentally determined, and modeling individual molecular motors is computationally intensive, we use a simplified approach that has been used previously to capture the effective overall force by motor proteins on each centrosome (13, 27).

We consider the following force-balance equation for the movement of centrosome *c* in the overdamped limit:

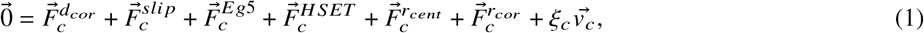

Where 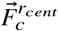 is a repulsive force between centrosomes. We solve a system of *c* equations for the velocity of each centrosome, 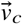, and use the velocity to determine the new location of each centrosome. Due to MT dynamics and stochastic force generation, a new set of forces in Eq. (1) are calculated at every time point, determining the corresponding centrosome velocity. The velocity is scaled by a constant drag coefficient, ξ, to account for the viscosity of the cytoplasm within the cell (Table S1) (56).

While the force by each motor population, dynein, Eg5, and HSET, is consistent on every MT they are bound to, we carefully consider how each force is felt by the centrosome center, and therefore contributes to centrosome movement. Stoke’s Law states that the drag on a spherical object is dependent on the viscosity of the fluid and the radius of the sphere when in free space. However, it is well established that the drag on a sphere increases when it is centered inside a confined spherical region (57). In this model, however, rather than a sphere, we have a centrosome with an attached radial array of MTs that are asymmetrically distributed and changing over time (Fig. S1 C,D in the Supporting Material). Our system is dynamic, with changing MT number, MT lengths, and centrosome position at every time step (Fig. S1 A in the Supporting Material). Studies have explored the drag on a symmetric and centered MT aster, where drag was an increasing function of MT volume fraction (49). However, they do not consider multiple asters, or how asters interact with each other. Further theoretical studies reveal that confinement and proximity to a boundary increases drag, but does not explore drag on non-solid objects (58). While these studies do not capture the effective drag on two asymmetric asters interacting with each other within a cell, they do provide insight into how forces should be scal ed to) account for this geometry. We therefore consider these changes to drag in an exponential damping term of the form 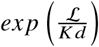 on each motor-dependent force, where ℒ defines a MT length scale, *d* defines the distance between the centrosome and the point of force application, and *K* is a constant parameter. Additional details are provided in the Supporting Material in Fig. S2 and Table S3 to show both model sensitivity and the scaling of this term with respect to ℒ.

### Cortical Forces

Dynein is a minus-end directed motor that is localized at the cell cortex (cor) during mitosis where it binds to MT plus-ends, generates a pulling force on the MT, and contributes to a net force that drives the centrosome closer to the boundary (9, 47, 48, 59–62), as illustrated in Fig. 1 C(iv). Cortical dynein is assumed to be uniformly distributed along the boundary and each MT plus-end within a distance 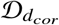 to the boundary has a probability 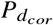 of binding to dynein. Following a standard Monte Carlo Method, we choose *n*_*d*_ from a uniform distribution, *n*_*d*_ ∈ 𝒰 [0, 1], and binding to dynein will occur if 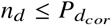. The pulling force generated by cortical dynein on the *i*^*th*^ MT nucleated from the *c*^*th*^ centrosome follows a standard linear force-velocity relationship (63):

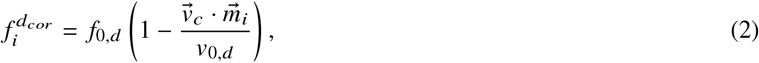

where *f*_0,*d*_ is the stall force of dynein, *v*_0,*d*_ is the walking velocity of dynein, 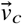 is the velocity of centrosome *c*, and 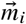 is the unit vector in the direction of MT *i*. The total pulling force by cortical dynein on the *c*^*th*^ centrosome in the direction of the *i*^*th*^ MT,

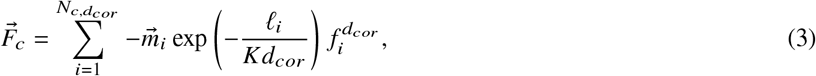

where 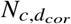 is the total number of MTs on centrosome *c* that bind to cortical dynein, *ℓ* _*i*_ is the length of MT *i, d*_*cor*_ is the minimal distance between centrosome *c* and the cell cortex, and *K* is a scaling factor. This force will pull the centrosome in the direction of 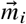, towards the cell cortex. MTs will stay bound to cortical dynein until the end of the MT is greater than a distance 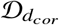 from the cell cortex, at which time it begins depolymerizing at velocity *v*_*s*_. As described earlier, the exponential term accounts for a higher drag on the centrosome due to MT length, density, and proximity to the cell boundary (Fig. S2 in the Supporting Material).

Alternatively, if the random number,, is greater than the probability of binding to dynein, 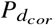, the MT instead continues to grow and slips along the boundary (47, 64) (Fig. 1 C(v)). For simplicity, we do not allow a MT to be bound to cortical dynein and grow/slip against the cortex simultaneously. The pushing force is described as:

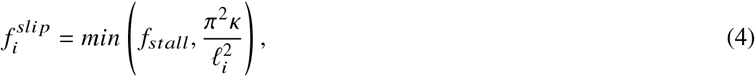

where *f*_*st all*_ is the stall force of a MT and *κ* is the bending rigidity of the MT. This force is also dependent on MT length, *ℓ*_*i*_, such that longer MTs are more likely to buckle than shorter MTs. The pushing force felt back on the *c*^*th*^ centrosome by *N*_*c,slip*_ MTs is then:

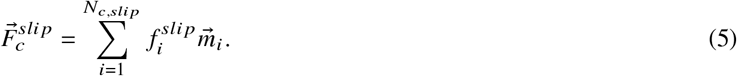

We note that this force already accounts for length-dependence, and long MTs are unlikely to generate significant force because they are more likely to buckle. Therefore, we do not consider the additional exponential scaling in forces derived by MT pushing against the cell boundary. Pushing MTs also experience a slight angle change of *θ* and the corresponding unit direction vector 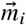 and angle *α*_*i*_ are then updated. A MT will stop pushing against the cell cortex if the end of the MT is greater than a distance 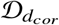 from the cell cortex. Alternatively, if 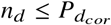 and the end of the MT is within 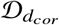 from the cell cortex, a pushing MT can then bind to cortical dynein.

### Interpolar Forces

Interpolar MTs can experience pushing or pulling forces by being bound to opposing MTs by either Kinesin-5 (Eg5, plus-end directed) or Kinesin-14 (HSET, minus-end directed), respectively (Fig. 1 (i),(iii)). Specifically, we define interpolar MTs as those having an angle within *π /*2 of the vector between the centrosomes (Fig. 2 A). Forces from Eg5 are necessary for centrosome separation early in mitosis, as loss of Eg5 prevents centrosome separation and results in monopolar spindles (6, 30, 65, 66).

HSET is localized along interpolar MTs and is involved in both antiparallel MT sliding and parallel MT bundling (7). However, since we do not explicitly model crosslinking activity by motors or passive crosslinker proteins, we consider only HSET activity on antiparallel MTs. HSET that is bound to antiparallel MTs is antagonistic to Eg5 and contributes to spindle maintenance during mitosis (32, 38, 67).

Interpolar MTs *i, j* nucleated from centrosomes *c, k*, that are within a distance 𝒟 _*Eg*5_ or 𝒟_*H SET*_ will have a probability of binding to Eg5 (*P*_*E*_) and/or HSET (*P*_*H*_) and generating force. Using a Monte Carlo Method, if a random number *n*_*E*_,*n*_*H*_ is less than *P*_*E*_, *P*_*H*_, binding of Eg5 and/or HSET occurs, respectively. The force on each MT by either Eg5 or HSET follows Eq. (2) with stall forces *f*_0, *Eg*5_, *f*_0, *H SET*_ and walking velocities *v*_0, *Eg*5_, *v*_0, *H SET*_, respectively. As MTs nucleated from both centrosomes are bound, we consider the net velocity of each centrosome in the force-velocity equation. The net velocity of centrosome *c* is therefore calculated as 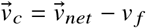 where 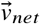 is the relative velocity between centrosomes *c* and *k*, and *v*_*f*_ is the poleward flux, the constant depolymerization of MT minus-ends on interpolar MTs (68, 69). The force felt on centrosome *c* due to Eg5 and HSET motor activity is

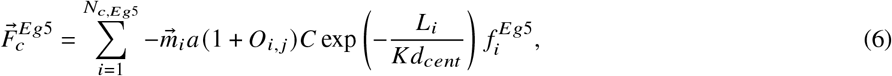

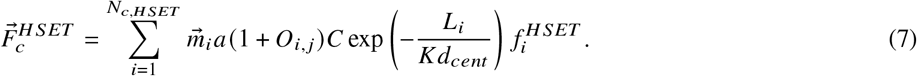

*N*_*c, Eg*5_ and *N*_*c, H SET*_ are the total number of MTs on centrosome *c* that are bound to Eg5 and HSET, respectively. *L*_*i*_is the distance between the centrosome *c* to the point where the motor binds to the *i*^*th*^ MT, *d*_*cent*_ is the distance between centrosomes *c* and *k*, and *C* is a constant scaling factor to account for both passive crosslinkers at antiparallel MT overlap regions (10, 12, 70, 71) and motor-dependent crosslinking activity by HSET and Eg5 (38, 40). The sensitivity of the model (defined by bipolar spindle length) to parameter *C* is shown in Table S3. If the angle of intersection between MTs *i* and *j, ϕ*_*i,j*_∈ [90°, 120°], then *a* = 1 and if *ϕ*_*i,j*_ *>* 120°, then *a* = 2; this allows interpolar MTs that are closer to antiparallel to generate more force as *ϕ*_*i,j*_ increases, simulating force by multiple motor proteins.*O* _*i,j*_ is the overlap distance of interpolar MTs *i* and *j* and is calculated as the minimum of, *ℓ*_*i*_, *ℓ*_*j*_ or *η*_*i,j*_, calculated as the law of cosines between the two MTs (Fig. 2 B). The equations scale with interpolar overlap distance *O*_*i,j*_ to account for force generation by multiple motors as this distance increases. For each interpolar interaction, the same equations are solved to calculate the force on centrosome *k*, using *L*_*j*_ in the exponential scaling term and 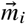, the unit direction vector of MT *j*, to determine the direction of the force. In Fig. S2 C in the Supporting Material we plot the relationship between *L*_*i*_ and the exponential scaling term, showing a decrease in force scaling with increased distance from the centrosome to motor-derived force.

### Spindle-Pole Dynein

In addition to its localization at the cell cortex, dynein is highly localized to spindle poles (sp) during mitosis (Fig. 1 (ii)), where it is necessary for the maintenance of MT minus-end focusing and spindle pole integrity (54, 72, 73). We allow MTs nucleated from opposing centrosomes to have a probability 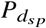 of binding to dynein anchored to MTs near centrosomes if they get within a distance 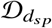 from the center of the centrosome. This motor-MT interaction is the same as Eq. (2). The force on centrosome *c* by dynein localized at spindle poles is calculated by

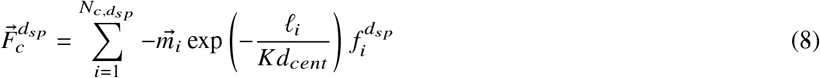

and is scaled to account for MT length, density, and proximity to the other centrosome. 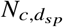 is the total number of MTs on centrosome *c* that bind to dynein localized at centrosome *k*. An equal and opposite force is felt on centrosome *k*.

### Repulsive Force

We consider a repulsive force between centrosomes to be activated if the distance between centrosomes, *d*_*cent*_, is less than 𝒟_*r*_ (Table S1). The force applied to centrosome *c* if this distance argument is achieved is:

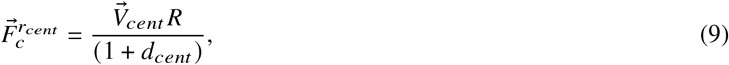

where 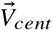 is the unit vector between centrosomes *c* and *k* (Fig. 2) and *R* is a scaling factor.

### Cell Culture, siRNA, Cell Line Generation

hTERT-immortaized Retinal Pigment Epithelial (RPE) cells were maintained in Dulbecco’s Modified Essential Medium (DMEM) supplemented with 10% fetal bovine serum (FBS) and 1% Penicillin and Streptomycin and maintained at 37°C with 5% CO_2_. Depletion of Nuf2 and Afadin was achieved by transient transfection of a pool of four siRNA constructs (Nuf2 target sequences: 5’-gaacgaguaaccacaauua-3’, 5’-uagcugagauugugauuca-3’, 5’-ggauugcaauaaaguucaa-3’, 5’-aaacgauagugcugcaaga-3’; Afadin target sequences: 5’-ugagaaaccucuaguugua-3’, 5’-ccaaaugguuuacaagaau-3’, 5’-guuaagggcccaagacaua-3’, 5’-acuugagcggcaucgaaua-3’; Dharmacon, Lafayette, CO) at 50 nM using RNAiMAX transfection reagent according to manufacturer’s instructions. Knockdown conditions were performed alongside a scrambled control (siScr) with a pool of four non-specific sequences (5’-ugguuuacaugucgacuaa-3’, 5’-ugguuuacauguuguguga-3’, 5’-ugguuuacauguuuucuga-3’, 5’-gguuuacauguuuuccua-3’; Dharmacon, Lafayette, CO). Depletion was confirmed by qPCR with primers for Nuf2 (F:5’-taccattcagcaatttagttact-3’, R:5’-tagaatatcagcagtctcaaag-3’ ;IDT, Coralville, IA), Afadin (F:5’-gtgggacagcattaccgaca-3’, R:5’tcatcggcttcaccattcc-3’; IDT, Coralville, IA), and GAPDH (F:5’-ctagctggcccgatttctcc-3’, R:5’-cgcccaatacgaccaaatcaga-3’; IDT, Coralville, IA) as a control. Nuf2 makes up one of the four arms of the Ndc80 complex, which attaches MTs to kinetochores, along with Hec1, Spc24, and Spc25 (74). Hec1 and Nuf2 dimerize in this complex, and knockdown of either protein destabilizes the other complex member, leading to loss of MT attachments to kinetochores (75). Therefore, knockdown of Nuf2 was further confirmed using immunofluorescence imaging with antibodies specific for Hec1 (Novus Biologicals, Littleton, CO) to assess kinetochore localization of the complex.

RPE cells stably expressing L304-EGFP-Tubulin (Addgene #64060, Watertown, MA) were generated by lentiviral transduction and placed under 10 *µ*g/mL Puromycin selection for 5-7 days. Expression of the tagged construct was confirmed by immunofluorescent imaging (76). RPE cells stably expressing GFP-centrin were previously described (77) and generously provided by Neil Ganem.

### Immunofluorescence Imaging

Cells were captured with a Zyla sCMOS (Oxford Instruments, Belfast, UK) camera mounted on a Nikon Ti-E microscope (Nikon, Tokyo, Japan). A 60x Plan Apo oil immersion objective was used for fixed-cell imaging and live-cell imaging of RPE cells expressing GFP-centrin to visualize centrosomes (78), and a 20x CFI Plan Fluor objective was used for live-cell imaging of RPE cells expressing GFP-tubulin (76).

### Fixed-cell Imaging and Analysis

Cells seeded onto glass coverslips were rinsed briefly in phosphate buffered saline (PBS) and placed in ice cold methanol for 10 minutes at −20°C. Coverslips were washed briefly with PBS and blocked in TBS-BSA (10 mM Tris at pH 7.5, 150 mM NaCl, 1% bovine serum albumin (BSA)) for 10 minutes. Cells were incubated with primary antibodies diluted in TBS-BSA (anti-*α*-tubulin (1:1500; Abcam ab18251, Cambridge, UK), anti-Ndc80 (1:500; Novus Biologicals, Littleton, CO), anti-Centrin (1:1000; Millipore 04-1624, Burlington, MA), anti-NuMA (1:150; Abcam ab109262, Cambridge, UK) for 1 hour in a humid chamber. Cells were washed in TBS-BSA for 10 minutes then incubated with fluorophore-conjugated secondary antibodies (Invitrogen, Carlsbad, CA) diluted 1:1000 in TBS-BSA + 0.2 *µ*g/mL DAPI for 45 minutes.

Fixed and live-cell image analysis was performed in NIS Elements. Fixed cell analysis of DNA area was quantified by gating a region of interest by DAPI fluorescence intensity. Spindle length was quantified by performing line scans along the long axis of the mitotic spindle and considering the spindle poles to be the two highest peaks in fluorescence intensity. Spindle morphology was characterized as bipolar, monopolar, or disorganized, where monopolar spindles were characterized by spindle length being less than half the average bipolar spindle length, and disorganized spindles had indistinguishable spindle poles. All analysis performed and all representative images are of a single focal plane. Background was subtracted by the rolling-ball algorithm (79) and contrast was adjusted in ImageJ to prepare fixed-cell images and GFP-centrin live-cell images for publication. Statistical analysis was performed in Excel; two-tailed Student’s *t*-test was used for comparisons between two groups.

### Live-cell Imaging

RPE cells stably expressing *α*-tubulin-EGFP were seeded onto a 6-well plate. NIS elements HCA jobs software was used to enable multi-coordinate, multi-well imaging in a single z-stack (0.67 *µ*m per pixel) (76). Images were captured every 5 minutes for 16 hours. Analysis was performed on at least 40 mitotic cells.

RPE cells stably expressing GFP-centrin were seeded onto glass coverslips and placed in a sealed chamber slide with 100 *µ*l of media. Single cells entering mitosis were captured at 60x in a single z-stack (0.11 *µ*m per pixel) every fifteen seconds for the duration of mitosis or until centrosomes were no longer in the same plane. Spindle length fluctuations were quantified as the average number of peaks per minute, rather than the total number of peaks per trace to account for changes in movie duration.

## RESULTS AND DISCUSSION

### Spindle formation occurs independently of stable microtubule interactions with chromosomes

To better define the extent to which stable end-on MT attachments to chromosomes are dispensable for bipolar spindle structure, we used immunofluorescence imaging approaches to observe cells depleted of Nuf2 (siNuf2), a protein essential for stable MT binding to kinetochores (Fig. 3 A). While it has been established that a bipolar spindle can form in the absence of stable MT attachments to kinetochores (33, 34), performing these experiments in house provides valuable data that can be used to inform and validate our model. We use RPE cells for mitotic analysis, which are a well characterized, diploid, immortalized mammalian cell line. We stained cells with DAPI to label chromatin and α-tubulin to label MTs. We used siRNA to specifically target Nuf2 for depletion and then assessed mitotic spindle structure. Consistent with previously described work (32–34), we find that Nuf2 depletion leads to a marked decrease in Hec1 localization at kinetochores and dispersion of chromosomes throughout the cell (Fig. 3 A,B). Additionally, Nuf2 depleted cells exhibit an increase in spindle length compared to the control condition (17 *µ*m for Nuf2 depletion and 14 *µ*m for control, Fig. 3 C). Together, this indicates the failure to form stable MT attachments to kinetochores following Nuf2 depletion. Despite these differences, spindle morphology remains largely bipolar in the Nuf2 depleted condition, with more than 90% of cells achieving bipolarity (Fig. 3 D). These data confirm that kinetochores and kinetochore-derived forces are not required for bipolar spindle formation and maintenance, validating our choice to omit chromosome-derived forces from our model.

### A biophysical model captures bipolar spindle formation and maintenance

To inform our model and validate model outputs, we performed live-cell imaging of RPE cells stably expressing an *α*-tubulin-EGFP transgene (Fig. 4 A) or a GFP-tagged centrosome marker (GFP-centrin) (Fig. 4 F,G). Spindle MTs are anchored at centrosomes by crosslinking and motor proteins to form spindle poles (54, 80), allowing analyses of either spindle pole or centrosome position to be used to quantify centrosome movement in space and time. We used RPE cells expressing *α*-tubulin-EGFP to inform initial conditions of the model (Fig. 4 A). We quantified intracentrosomal distances just prior to nuclear envelope breakdown (NEB), defined as the first point in time at which GFP-tubulin is no longer visibly excluded from the nuclear region. This analysis reveals a wide distribution, with initial centrosome distances ranging between 3.9 and 16.6 *µ*m (Fig. 4 C). To mirror this distribution of centrosome positions in our model, we initialize centrosomes to be randomly placed at least 7.5 *µ*m from the center of the cell, achieving a range of distances between 4.2 and 14.75 *µ*m (Fig. 4 C).

**Figure 4:**
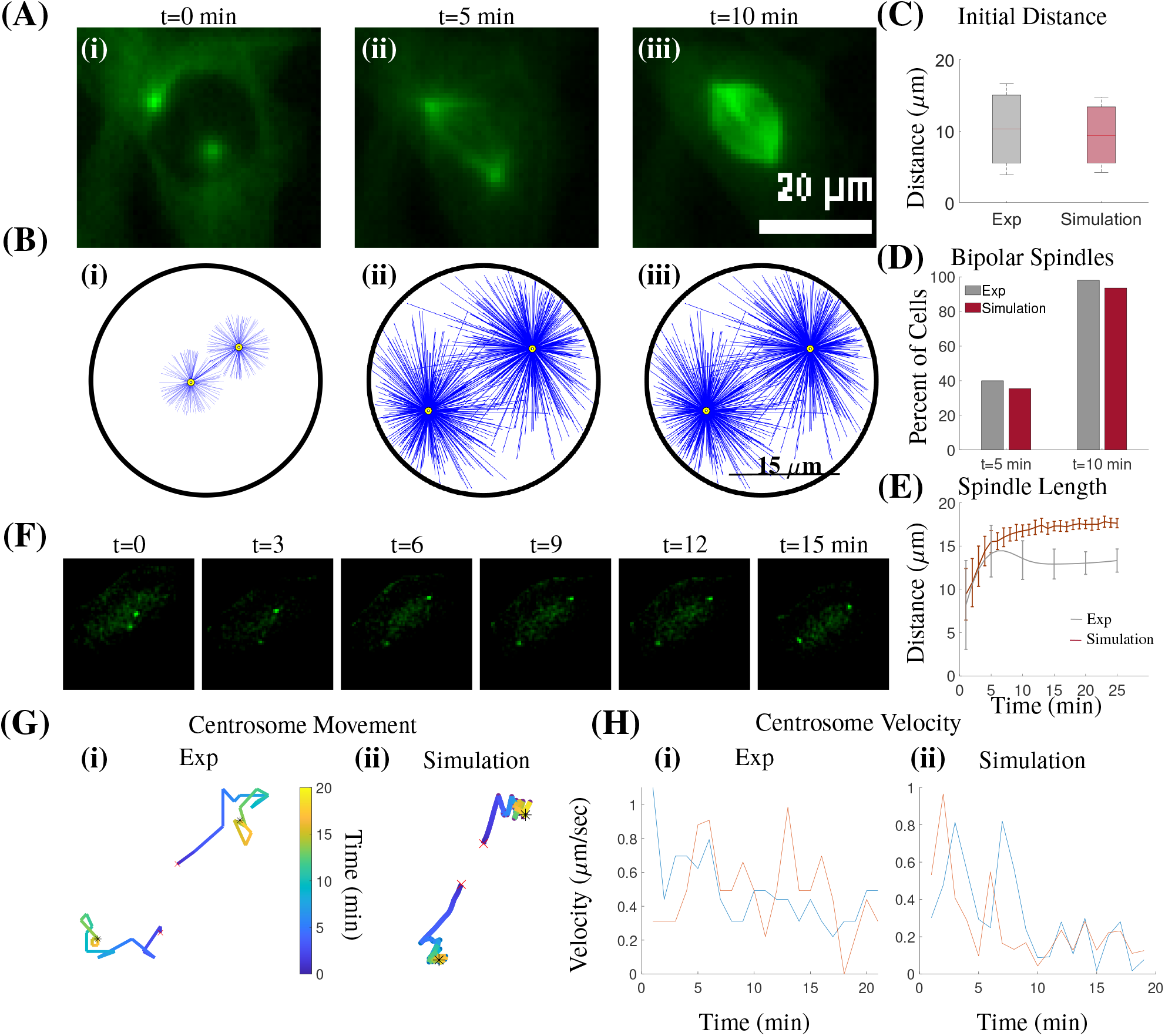
Stochastic force-balance model captures centrosome movement and bipolar spindle formation. (A) Still frames from live-cell imaging of RPE cells expressing tubulin-EGFP from the time point before nuclear envelope breakdown (NEB) at t=0 min in (i) to spindle bipolarity at t=10 min in (iii). Image acquisition parameters for live-cell analysis limit detection to more stable populations of MTs, primarily kinetochore MT bundles. However, fixed cell analyses confirm astral and interpolar MT populations are also prevalent in these same conditions (see also Fig. 3 A). (B) Still frames from a single simulation showing initial centrosome positioning at t=0 min in (i) to spindle bipolarity at t=10 min in (iii). The corresponding simulation is shown in Movie M1 in the Supporting Material. (C) Distributions of initial distance between spindle poles from live-cell imaging (Exp) and simulations. (D) Plot of the fraction of cells (Exp) and simulations that achieve bipolarity by 5 and 10 min. (E) Plot of the spindle length over time from live-cell imaging (Exp) and simulations. Error bars are standard deviation. Biological data are captured at 5 min increments; a cubic spline is used to generate the curve. All averages for (C)-(E) calculated from at least 40 cells and 30 simulations. (F) Still frames from live-cell imaging of RPE cells expressing GFP-centrin. (G) (i) Experimental traces of centrosome movement from cell shown in (F), where color denotes time (min). (G) (ii) Traces of centrosome movement from a single simulation, where the two lines correspond to the two centrosomes, and color denotes time (min). Red ‘x’ is initial centrosome position, black asterisk is final centrosome position. (H) (i) Centrosome velocities over time from cell shown in (F, G (i)). (ii) Centrosome velocities over time from simulation shown in (G)(ii). Each line is a centrosome.

Live-cell imaging was used to monitor centrosome movement and spindle bipolarity, capturing centrosome separation at early time points (Fig. 4 E,F,G(i)) until an eventual bipolar spindle is achieved and maintained at an average spindle length of 12 *µ*m (Fig. 4 E). Imaging analyses further reveal that 40% of cells achieve spindle bipolarity (spindle length *>*10*µ*m) by 5 min and 96% by 10 min (Fig. 4 D). Quantification of bipolar spindle length from live-cell imaging is consistent with fixed-cell image analysis of RPE cells with stable MT-chromosome attachments in Fig. 3 C (siScr). By tracking individual centrosome positions in time, we calculate that they have a velocity less than 1 *µ*m/sec. While mitotic progression has been well characterized, performing this analysis provides data to directly integrate and compare with our model.

We have parameterized our model such that mitotic timing, bipolar spindle length, and centrosome velocity closely match our experimental measurements. Where available we used parameters that have been well established (Table S1), and where necessary we have defined and optimized new parameters to closely capture biological phenomena (Tables S1, S3, S4). Late time points of our model resemble a bipolar spindle with asymmetrically distributed MTs, with an increased density towards the center of the spindle structure (Fig. 4 B(iii), Fig. S1 D). MTs in the interpolar region (the region between spindle poles) are interacting and generating force, allowing the maintenance of this bipolar configuration (Figs. 5, 6). Model analysis shows that 35% of simulations achieve spindle bipolarity by 5 min and 94% by 10 min (Fig. 4 D, bipolar defined as having a spindle length ≥1/ 2 of the final average spindle length). Furthermore, an average bipolar spindle length of 17 *µ*m is achieved (Fig. 4 E). While this is a longer spindle length than that seen in control RPE cells (Fig. 3 C (siScr)), it is consistent with measured spindle lengths from RPE cells depleted of Nuf2 (Fig. 3 C (siNuf2)) which, like our model, lack kinetochore-derived forces. Centrosome movement and velocity similarly resembles biological results (Fig. 4 G,H), suggesting that our parameterized model closely captures the dynamics of mitotic progression. We use this as our model base case throughout this work.

**Figure 5:**
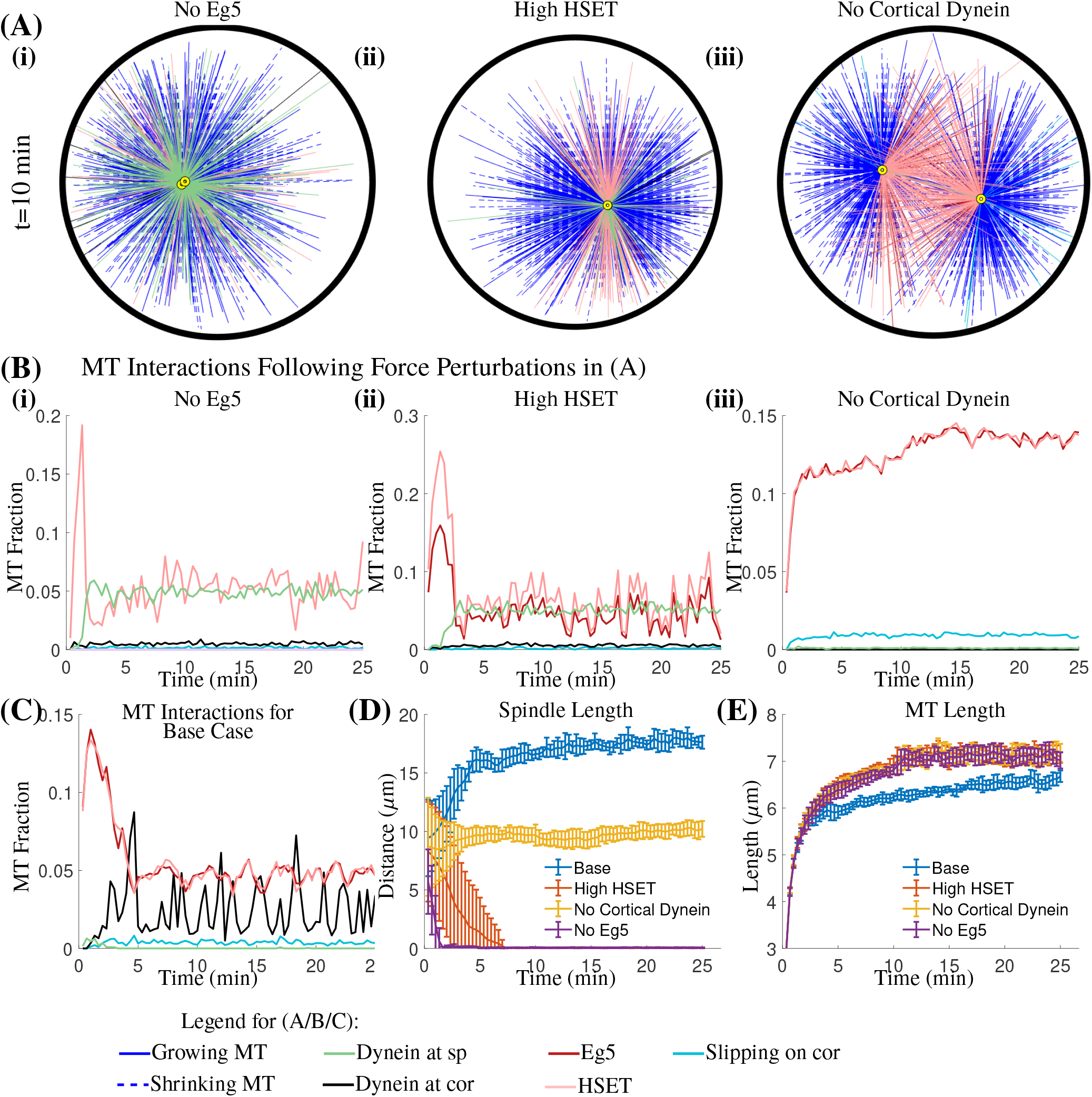
Force perturbations impact spindle bipolarity. (A) Still frame from a simulation at t=10 min with: (i) no Eg5 binding, (ii) high HSET binding, or (iii) no cortical dynein binding. Colors indicate the force generated by each MT interaction, defined in the legend. (B) Plot of the average percent of MTs in each force-generating state over time with: (i) no Eg5 binding, (ii) high HSET binding, or (iii) no cortical dynein binding, and (C) the base case. (D) Plot of the average distance between spindle poles over time for the base case and each single force perturbation. (E) Plot of the average length of MTs over time. All averages are of 10 simulations and error bars shown correspond to standard deviation.

**Figure 6:**
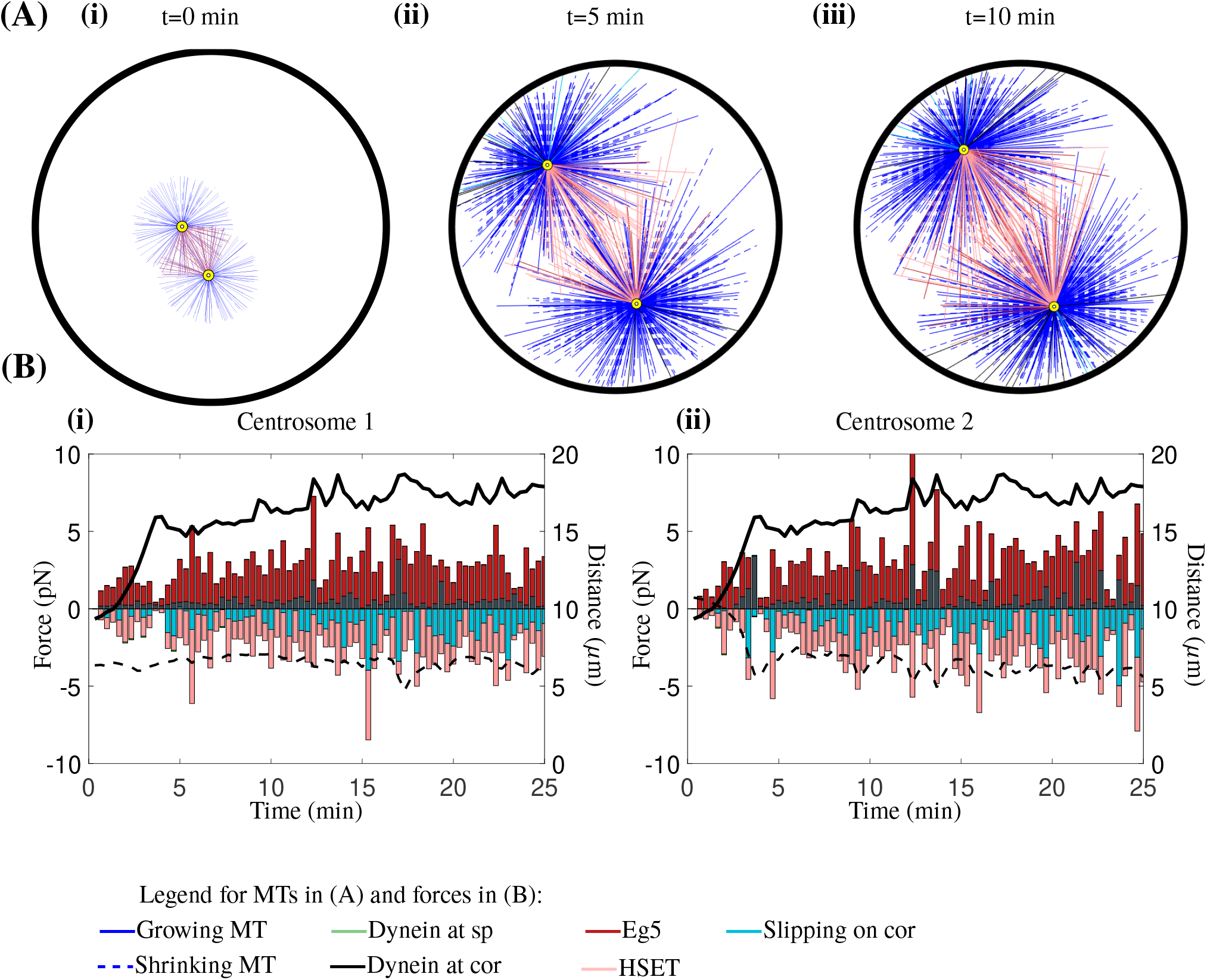
Forces are dynamic over time, enabling the formation and maintenance of a bipolar spindle. (A) Still frames from a simulation from initial centrosome positioning (i) to spindle bipolarity (iii). MT color represents its “state”, defining the force that it generates. (B) Force plots of centrosome “1” (i) and centrosome “2” (ii) in the direction of the vector between the two centrosomes, where a positive force brings centrosomes together and negative force pushes centrosomes apart. Black solid line shows spindle length over time and black dashed line shows the absolute minimum centrosome distance to the cell cortex over time.

### Motor protein perturbations alter spindle bipolarity

The mitotic spindle has been extensively studied, and our understanding of the force requirements for spindle bipolarity has been determined primarily through experimental manipulation of force-generating motor proteins. While informative, biological assays can induce potential off-target effects and can impact multiple cellular processes. In contrast, mathematical and computational modeling allows for the specific modulation of individual motor populations and affords temporal control of such perturbations to defined stages of mitosis. Therefore, to determine how motor proteins considered in our model impact spindle bipolarity, we independently perturbed motor function of Eg5, HSET, and cortical dynein. We accurately reflect perturbed motor activity by altering the binding probability of the motor *β* from the base case of *P*_*β*_ = 0.5. (Table S1). All other parameters remain unchanged from the base case, allowing us to specifically determine the impact of altered motor activity on spindle bipolarity.

Biological data indicates that loss of Eg5 activity results in spindle pole collapse and the formation of a monopolar spindle (6, 30, 65–67). To determine if our model is able to capture this phenomenon, we simulate loss of Eg5 activity by setting the probability of Eg5 binding to MTs (*P*_*E*_) to zero. Our simulations with loss of Eg5 activity result in failure to establish a bipolar spindle (Fig. 5 A(i),D), and maintained spindle collapse through the duration of the simulation. Consistent with the requirement of Eg5 for centrosome separation and early bipolar spindle formation in cells, our simulations with no Eg5 activity show that centrosomes collapse to a monopolar spindle is immediate, with a monopolar spindle being formed in less than 2 min (Fig. 5 D). Analysis of the fraction of MTs bound to motor proteins over time reveals that HSET activity remains unchanged from the base condition (Fig. 5 B(i),C). However, spindle pole-localized dynein becomes relevant with loss of Eg5, where it helps to maintain close proximity of centrosomes following spindle pole collapse (Fig. 5 B(i),C).

Biological results also show that high HSET activity increases the frequency of monopolar spindles (81–83). To test that our model accurately reflects this role of HSET activity, we mimic HSET overexpression by setting the probability of binding to MTs (*P*_*H*_) equal to one. Consistent with published biological data, our model captures monopolar spindle formation with high HSET activity (Fig. 5 A(ii),D). We observe that monopolar spindle formation occurs almost immediately, with all simulations having a fully collapsed spindle by t=5 min (Fig. 5 D). Similar to the condition with no Eg5, the fraction of MTs bound to spindle pole-localized dynein is increased with high HSET activity compared to the base condition (Fig. 5 B(ii),C). These results suggest that spindle pole dynein is similarly important in maintaining a monopolar spindle when HSET activity is high.

Due to the multiple functions of dynein at spindle poles, kinetochores, and the cell cortex (54, 59, 60, 72), biological approaches have been unable to discern the specific role of cortical dynein in bipolar spindle formation. To address this limitation, cortical dynein activity was depleted in our model by setting the probability of binding to MTs 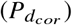 to zero. Our simulations indicate that specific loss of cortical dynein results in shorter bipolar spindles, decreased from 17 *µ*m in the base case to 10 *µ*m (Fig. 5 A(iii),D and Movie M2). We additionally see a greater than 2-fold increase in MTs bound to Eg5 and/or HSET when cortical dynein activity is absent compared to the base case, where the percent of MTs bound to both Eg5 and HSET increases from 6% in the base case to 15% in the absence of cortical dynein (Fig. 5 B(iii),C).

None of the single motor protein perturbations described have a significant impact on average MT length compared to the base condition (Fig. 5 E). As such, the changes in bipolar spindle length following perturbations to motor activity are strictly a result of altered forces on the centrosomes and not a consequence of limitations imposed by altered MT lengths. Combined, these results indicate that our model both captures known changes in bipolar spindle length following loss of Eg5 or overexpression of HSET, and demonstrates a decrease in steady-state spindle length following loss of cortical dynein.

### Cortical dynein is a primary regulator of bipolar spindle length

The biophysical model used here to describe and explore the dynamics of bipolar spindle formation and maintenance has the benefit of discretely defined MTs, each of which can generate force depending on its length and position relative to other intracellular components (Fig. 6 A, Movie M1). To explore how the magnitude and direction of forces on centrosomes change during spindle formation, we assessed each component of the force over time, with respect to 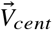, the unit vector between centrosomes (Fig. 2 A) (using the projection of the total forces in the direction of 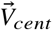). We considered a positive force to be one that increases spindle length (i.e. Eg5/cortical dynein) and a negative force to be one that decreases spindle length (i.e. HSET/pushing on the cell cortex/dynein at spindle poles).

To visualize how forces contribute to spindle dynamics, force plots for each centrosome were overlaid with curves for spindle length and the minimal centrosome distance to the cell cortex over time (Fig. 6 B(i)-(ii)). In our base case, where we have no perturbed motor activity, we find dynamic and reproducible force-dependent changes in spindle length. Our analysis shows that forces driving centrosome movement are dominated by Eg5 at early time points (t*<*5 min), consistent with the known biological role of Eg5 in mitosis (84–86). While averaging over many simulations of the base case show that a stable bipolar spindle length of 17*µ*m is achieved (Fig. 4 E, Table S2), analysis of a single simulation indicates that this is a quasi steady-state, where fluctuations in bipolar spindle length occur. Observing how forces change over time reveals that these fluctuations coincide with increased cortical dynein-derived force (Fig. 6 B). These data implicate cortical dynein in orchestrating dynamic changes to bipolar spindle length during mitosis.

### Cortical dynein drives fluctuations in spindle length after spindle bipolarity is achieved

To define the forces required for fluctuations in bipolar spindle length we first explored the consequences of perturbing cortical dynein pulling forces. To mimic loss of cortical dynein activity, we altered 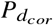, the probability of MTs binding to dynein at the cell cortex. As 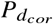 is reduced, bipolar spindle length decreases (17.9 *µ*m when 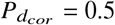 to 15.6 *µ*m when 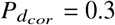, and 10.3 *µ*m when 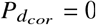) (Fig. 7 A, Movie M2), implicating cortical dynein in the regulation of steady-state bipolar spindle length.

**Figure 7:**
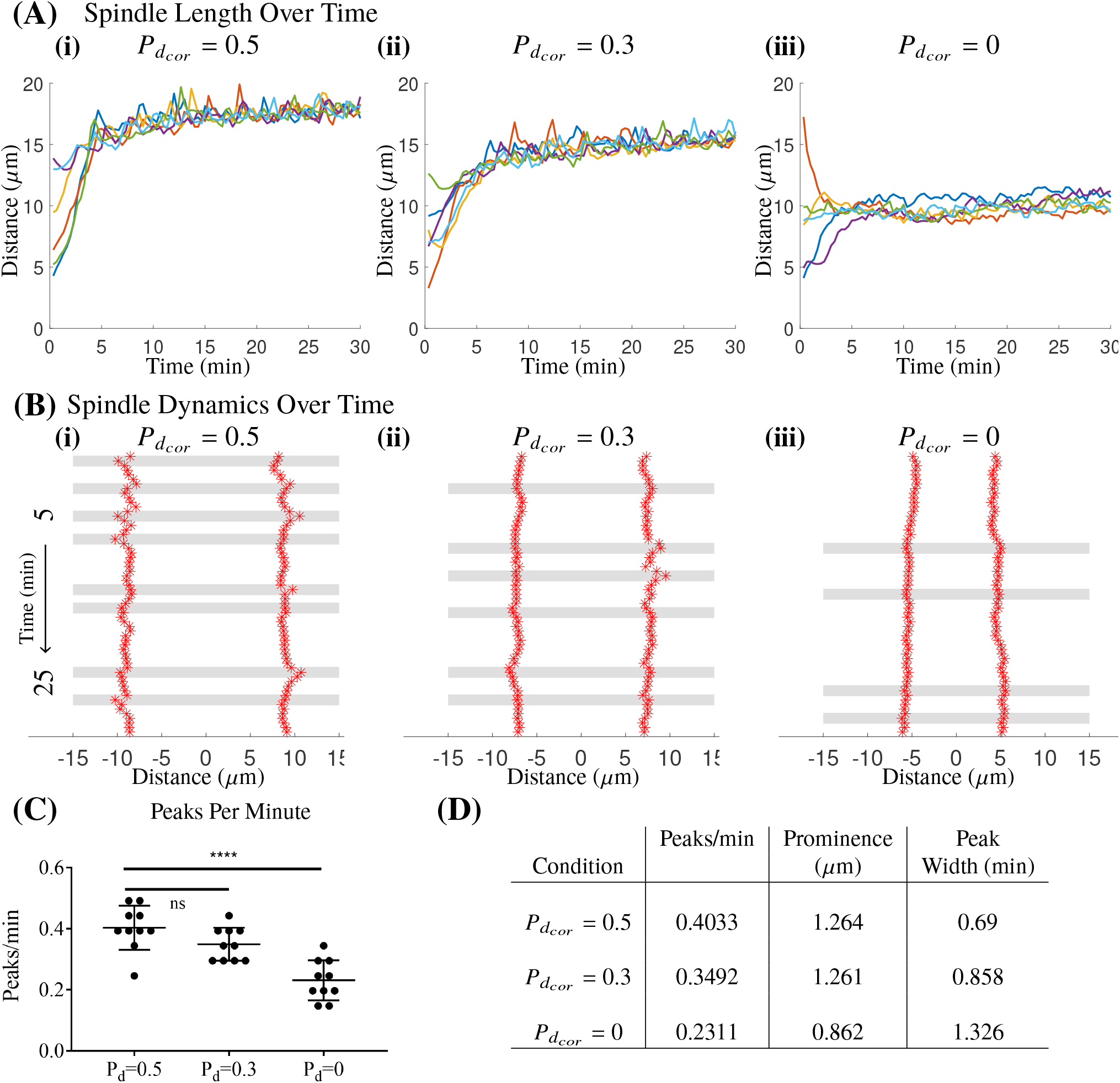
Cortical dynein regulates spindle length and bipolar spindle dynamics. (A) Curves of spindle length over time for 6 simulations with a dynein binding probability, 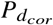, of 0.5 in (i), 0.3 in (ii), and 0 in (iii). (B) Representative kymograph of a single simulation with dynein binding probability 0.5 in (i), 0.3 in (ii), and 0 in (iii) from 5 to 25 minutes. Red asterisks are centrosome position plotted every 20 seconds, gray bars indicate prominent peaks in spindle length. (C) Plot of the number of peaks per minute from 10 simulations with dynein binding probability 0.5, 0.3, and 0. Each dot is a simulation, error bars are mean and SD. ****p*<*0.0001, ns indicates not significant. (D) Table showing average number of peaks per minute, average peak prominence, and average peak width over 10 simulations for each condition with dynein binding probability 0.5, 0.3, or 0.

To define a time-dependent relationship between bipolar spindle length and cortical dynein binding and pulling forces, we performed quantitative time-series analyses. The data is represented as a kymograph, a graphical representation of position over time, where the *y*-axis represents time (Fig. 7 B). In each plot, *x* = 0 is the center of the cell and *x* = −15, *x* = 15 are the cell boundaries. Red asterisks indicate centrosome position at 20 sec time intervals. We used peak prominence (87), defined as the vertical distance between the height of a peak and its lowest contour line, as a readout of significant changes in spindle length. Peaks identified as significant had a prominence greater than the minimum average standard deviation within spindle length traces between the conditions 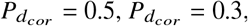 and 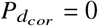. As shown in Fig. 7 B, C, D, we find that fluctuations in bipolar spindle length have both decreased frequency (peaks/min), decreased amplitude (prominence), and increased duration (width) when cortical dynein activity is decreased. Specifically, we see a 14% and 43% decrease in the number of peaks per minute from the base condition when 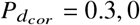, respectively. Furthermore, we see a 32% decrease in peak prominence when 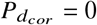, although we see no change when 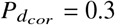, and a 24% and 92% increase in peak width when 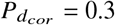, 0, respectively. Together, this data suggests that reduced cortical dynein activity alters the frequency, amplitude, and duration of bipolar spindle length fluctuations.

To determine if cortical dynein activity similarly impacts bipolar spindle length in cells, we performed fixed-cell imaging and analysis of pole-to-pole distance in RPE cells. We disrupted cortical dynein activity with either short-term chemical inhibition (Dynarrestin) or via depletion of Afadin, a protein involved in localizing dynein-NuMA complexes to the cell cortex (88, 89). Duration and concentration of Dynarrestin treatment was optimized to preferentially impair cortical dynein activity as previously described (89). Afadin depletion was validated by qPCR and confirmed by quantification of reduced cortical NuMA staining intensity (Fig. S3 in Supporting Material). Consistent with our modeling results, fixed-cell imaging reveals that average bipolar spindle length is reduced from 13 *µ*m to 11.1 *µ*m and 9.6 *µ*m in Nuf2 depleted cells following disruption of cortical dynein activity by Afadin depletion or Dynarrestin treatment, respectively (Fig. 8 A,B, Fig. S4 in Supporting Material). Similar results were observed in control cells with functional kinetochore attachments following treatment with Dynarrestin, with a reduction from 11.05 *µ*m to 8.9 *µ*m (Fig. S4 A,B in Supporting Material). While spindle length with Afadin depletion alone remains comparable to the control (siScr), depletion of Afadin in the absence of Nuf2 shows a decrease in spindle length that is not statistically different than what is seen following Dynarrestin treatment (Fig. 8 B), (Fig. S4 A,B in Supporting Material). These data raise the possibility that kinetochore MT attachments may stabilize spindle length in the absence of Afadin, thereby limiting the impact of decreased cortical dynein on bipolar spindle length.

**Figure 8:**
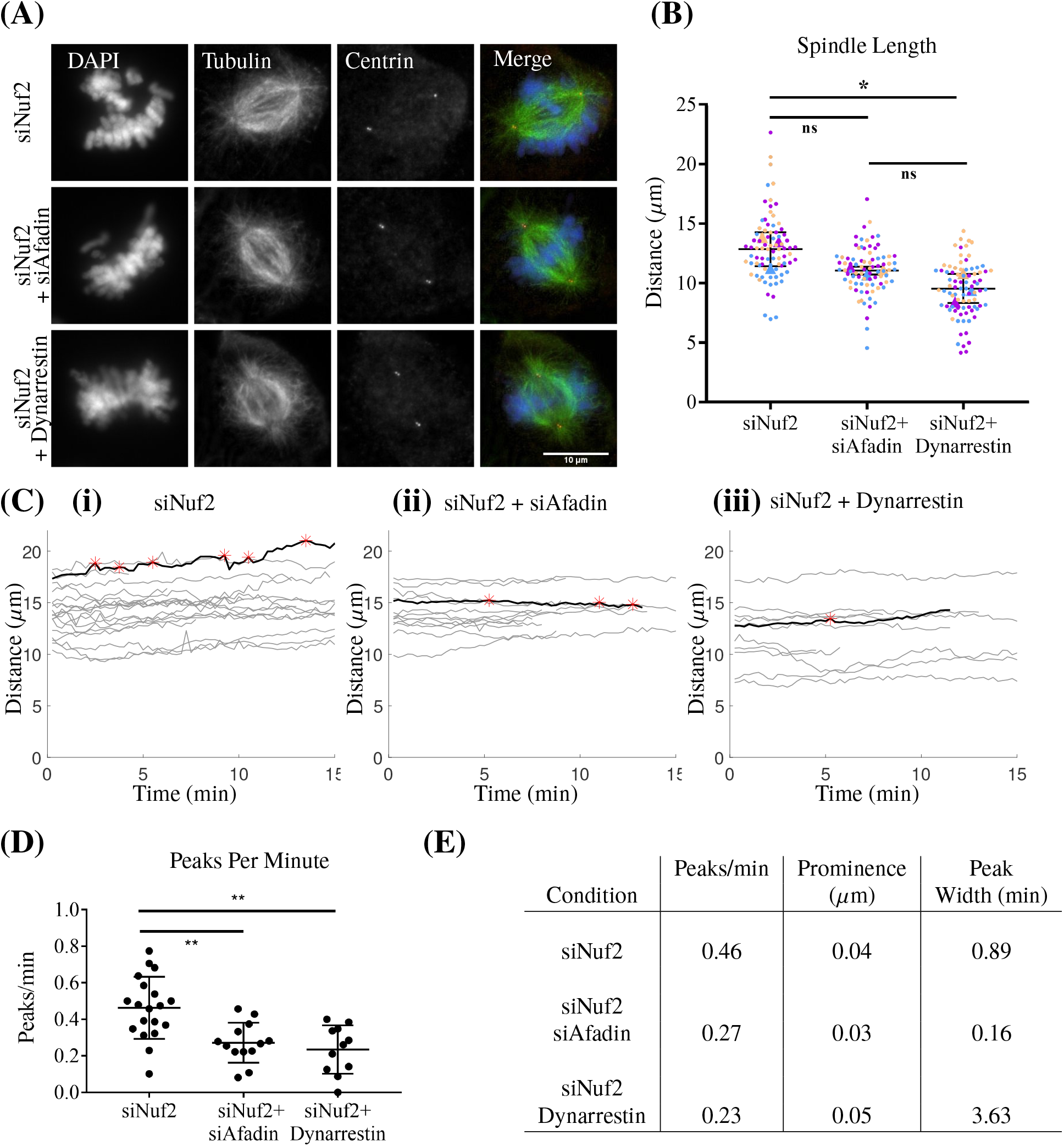
Fixed and live-cell imaging captures dynein-dependent changes in bipolar spindle length and spindle dynamics. (A) Fixed cell imaging of RPE cells stained for DAPI (DNA), Tubulin (MTs), and Centrin (centrosomes) in siNuf2, siNuf2+siAfadin, and siNuf2+Dynarrestin conditions. (B) Quantification of bipolar spindle length in siNuf2, siNuf2+siAfadin, and siNuf2+Dynarrestin conditions. Quantification performed on at least 25 cells from each condition for 3 biological replicates. Each color indicates a replicate and the average for each replicate is represented by a triangle of the same color. (C) Traces of spindle length over time of individual RPE cells expressing a GFP-centrin tag for siNuf2 (i), siNuf2+siAfadin (ii), and siNuf2+Dynarresetin (iii) conditions. Red asterisks represent significant peaks on the curve shown in black. (D) Quantification of the average number of peaks per minute in siNuf2, siNuf2+siAfadin, and siNuf2+Dynarrestin conditions. Significance determined by one-way ANOVA. (E) Table showing the average number of peaks per minute, the average peak prominence, and average peak width from each condition. At least 10 cells were captured and quantified for each condition. All error bars are SD. *p*<*0.05, **p*<*0.01, ns indicates not significant.

To test whether fluctuations in bipolar spindle length could be observed in cells, we next performed live-cell imaging of RPE cells expressing GFP-centrin (Movie M3). Similar to the analysis performed on simulations, significant peaks were determined by peak prominence. Prominent peaks were considered as those having a prominence greater than the minimum average standard deviation within spindle length traces from all six conditions (siScr, siAfadin, Dynarrestin, siNuf2, siNuf2+siAfadin, siNuf2+Dynarrestin). Consistent with our simulations, we observe an average of 0.36 peaks/min in control cells (siScr) and 0.46 peaks/min in cells depleted of Nuf2 (siNuf2) that lack stable chromosome attachments (Fig. 8 C,D,E, Fig. S4 C,D,E in the Supporting Material). We used Afadin depletion (siAfadin) or Dynarrestin treatment, as described previously, to determine if loss of cortical dynein activity impacts spindle length fluctuations. In Nuf2 depleted cells, we see a significant 41% and 50% decrease in the average number of peaks per minute with Afadin knockdown and Dynarrestin treatment, respectively (Fig. 8 C,D,E). We also see a significant 44% decrease in the number of peaks per minute in the absence of Afadin alone (Fig. S4C,D,E). These results are consistent with our model, where loss of cortical dynein decreases the number of peaks per minute by 43% (Fig. 7 C,D). However, we do not see a significant decrease with Dynarrestin treatment, indicating a possibility of altered spindle structure or stability following extended treatment through the duration of imaging. Together, model predictions and biological results implicate cortical dynein activity in spindle length fluctuations during mitosis.

### Modeling reveals that high Eg5 activity rescues spindle length fluctuations in the absence of cortical dynein

To further define the relationship between MT-derived forces and the maintenance of spindle bipolarity in the presence or absence of cortical dynein, we increased Eg5 activity thereby increasing the outward force on each centrosome (i.e. pushing away from each other). We find that increasing Eg5 activity, by increasing the binding probability of Eg5 to MTs (*P*_*E*_), significantly increases bipolar spindle length, regardless of cortical dynein activity (Fig. 9 A). However, reduced spindle length seen in the absence of cortical dynein is not restored with high Eg5 activity (Fig. 9 A(1),(6)), suggesting that cortical dynein pulling force, independent of Eg5 activity, is important in establishing and maintaining bipolar spindle length.

**Figure 9:**
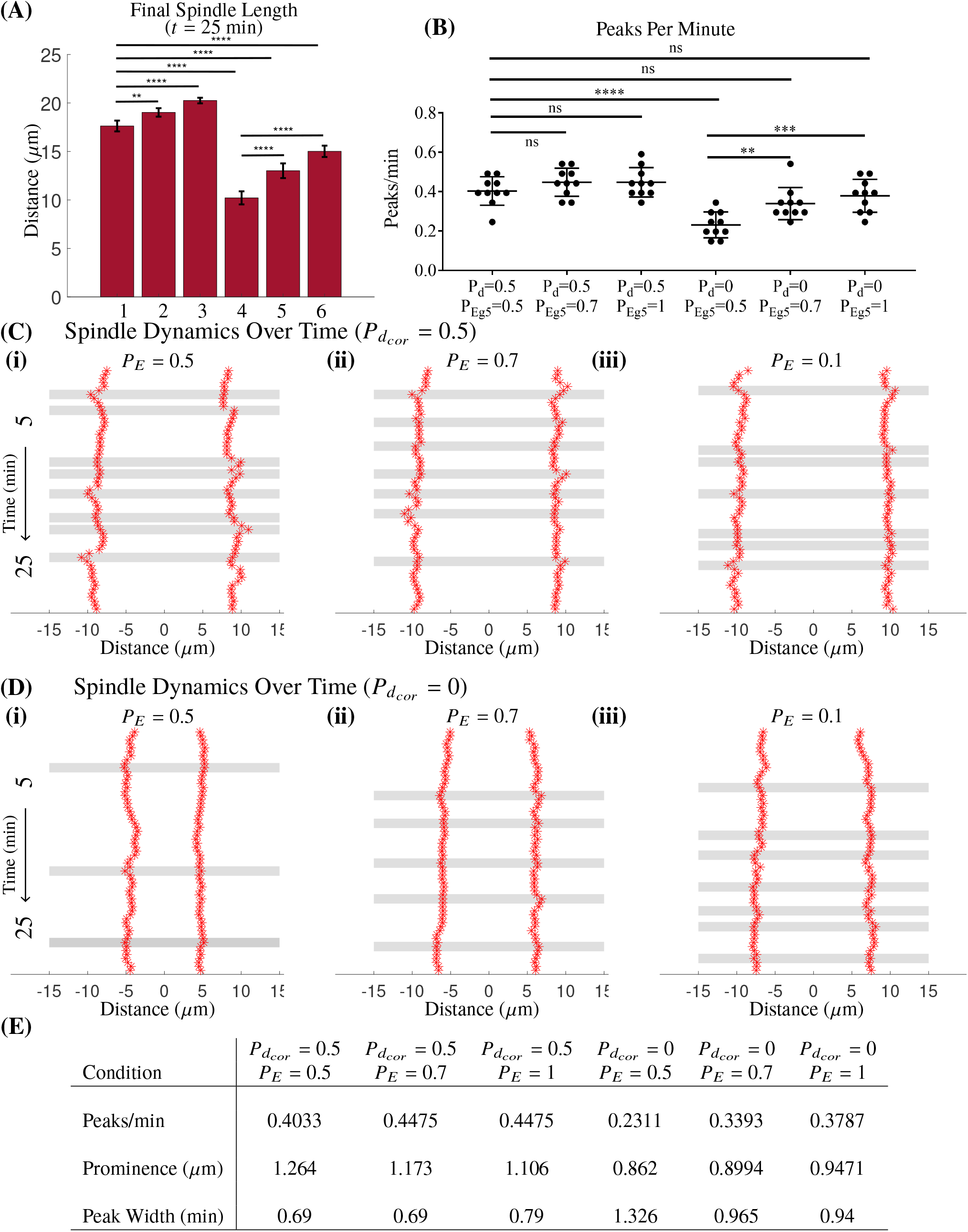
High Eg5 activity rescues spindle length fluctuations, but not steady-state spindle length in the absence of cortical dynein. (A) Bar graph of final spindle length from 10 simulations for each condition: (1) 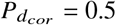, *P*_*E*_ = 0.5, (2) 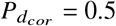, *P*_*E*_ = 0.7, (3) 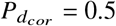, *P*_*E*_ = 1, (4) 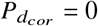, *P*_*E*_ = 0.5, (5) 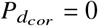, *P*_*E*_ = 0.7, (6) 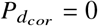, *P*_*E*_ = 1. (B) Quantification of the number of peaks per minute from 10 simulations for each condition Each dot is a simulation. (C/D) Representative kymographs of varied Eg5 binding probabilities, in the presence (C) or absence (D) of cortical dynein. Gray bars indicate prominent peaks in spindle length. (E) Table of the average number of prominent peaks, the average peak prominence, and the average peak width for each condition. All data averaged over 10 simulations for each condition. Error bars are mean and SD, **p*<*0.01, ***p*<*0.001, ****p*<*0.0001, ns indicates not significant.

To determine if Eg5 activity impacts spindle fluctuations in bipolar spindle length, we quantified the number of peaks per minute in simulations with increased Eg5 activity with and without cortical dynein. We find that in simulations with cortical dynein activity, increased Eg5 activity, either at intermediate (*P*_*E*_ = 0.7) or high (*P*_*E*_ = 1) levels, does not significantly impact the number of peaks per minute (Fig. 9 B,C,E, Movie M4). However, in the absence of cortical dynein activity, increased Eg5 activity rescues spindle length fluctuations to levels that are not significantly varied from the base condition (Fig. 9 B,D,E). Increased Eg5 activity does not, however, restore reduced peak prominence nor increased peak width in the absence of cortical dynein (Fig. 9 E). These results suggest that Eg5 activity cooperates with cortical dynein-derived forces to maintain fluctuations in bipolar spindle length.

### Cortical dynein is required for spindle bipolarity when HSET activity is high

HSET overexpression is prominent in many cancer contexts where its expression corresponds with increased cell proliferation (90, 91). This relationship with proliferation is independent of centrosome number, though in cancer contexts where centrosome number is amplified, HSET is additionally required to cluster extra spindle poles into a bipolar spindle (78, 92–96). We have confirmed that our model captures spindle-pole collapse in the context of high HSET activity (Fig. 5 A (ii), D). We then sought to further understand the sensitivity of spindle bipolarity to HSET activity. To test this, we incrementally increased the HSET binding probability in our model from its base level of *P*_*H*_ = 0.5. Our simulations indicate that spindle bipolarity is sensitive to HSET activity, such that the incidence of spindle pole collapse increases with high HSET activity, with only 40% of simulations forming a bipolar spindle when *P*_*H*_ = 0.8 and 0% when *P*_*H*_ = 0.9 or *P*_*H*_ = 1 (Fig. 10 A). To determine the force requirements for bipolar spindle formation in the presence of high HSET (*P*_*H*_ = 0.8), we explored a range of increasing cortical dynein activity and found that spindle bipolarity is rescued by cortical dynein activity in a concentration-dependent manner, with 90% of simulations forming a bipolar spindle when 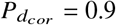 or 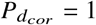 (Fig. 10 B).

**Figure 10:**
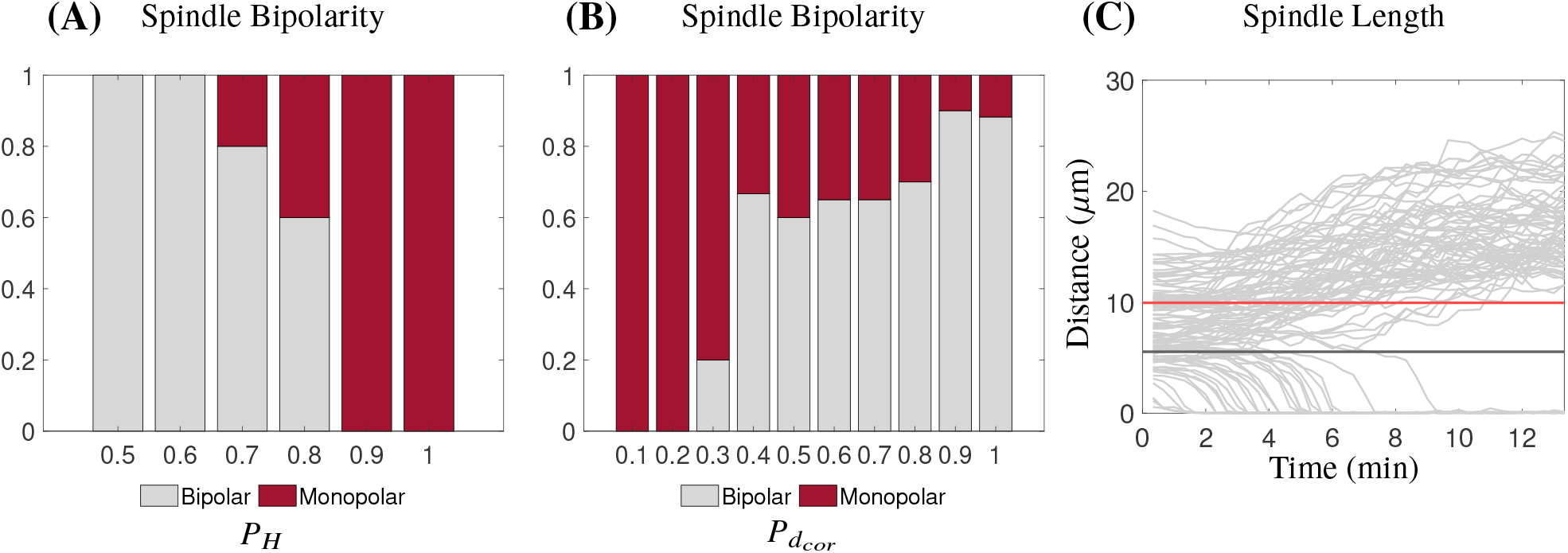
High cortical dynein promotes spindle bipolarity in the presence of high HSET. (A) Fraction of simulations that form a bipolar spindle with varying levels of HSET 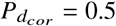. (B) Fraction of simulations that form a bipolar spindle in the presence of high HSET (*P*_*H*_ = 0.8) with varying levels of cortical dynein. (C) Plots of spindle length over time of simulations with high HSET (*P*_*H*_ = 0.8) and varying levels of cortical dynein that have simulations that form a bipolar spindle 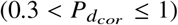. Red line is the average initial distance of centrosomes that separate (9.95 *µ*m) and black line is the average initial distance of centrosomes that collapse (5.4 *µ*m). Data from 20 simulations for each condition.

Work from other groups indicates that HSET-dependent motor activity is a dominant force in centrosome clustering once centrosomes reach a critical distance of 7-8 *µ*m from each other, whereas centrosome pairs are not impacted by HSET activity when they are 11-12 *µ*m apart (97). Consistent with this, we find that centrosomes collapse when they are, on average, initially 5.4 *µ*m apart, and instead form a bipolar spindle when initial centrosome distance is, on average, 9.95 *µ*m apart. Together, these results indicate that high cortical dynein activity and/or a large initial centrosome distance promotes bipolar spindle formation in the presence of high HSET.

## CONCLUSION

The biophysical model presented here forms and maintains a bipolar mitotic spindle through the balance of five MT-derived forces, including MT interactions with three key motor proteins: kinesin-14 (HSET), kinesin-5 (Eg5), and dynein (Figs. 1, 4, 6, Movie M1). While members of the kinesin-4, kinesin-6, and kinesin-8 families have been described as having roles in mitotic progression, their primary roles involve either chromosome condensation and alignment, or spindle midzone stability during anaphase (98–109). Since we are not explicitly modeling chromosomes, chromosome-derived forces, or anaphase spindle elongation, force contributions by these proteins were omitted. Our model was based on and validated using our experimental data defining centrosome position and time-dependent changes in spindle bipolarity in mammalian cells (Figs. 3, 4, 8).

Biological inhibition or knockdown of Eg5, and overexpression of HSET are shown to alter spindle bipolarity (6, 30, 65, 66, 81– 83). We manipulate motor activity in the model by perturbing the motor-MT binding probability, which accurately captures spindle collapse with no Eg5 or HSET activity (Fig. 5). To further inform the force balance between motor-derived forces through mitotic progression, future work could explore the spatiotemporal distribution of motor activity along MTs at the interpolar overlap regions. Simulating discretely localized proteins throughout the spindle structure would provide estimates for the required motor concentrations for spindle formation and maintenance.

The role of dynein activity in spindle formation and maintenance has been difficult to discern due to its localization and function at spindle poles, kinetochores, and the cell cortex during mitosis (54, 59, 60, 72). By defining each of these motor populations independently in our model, we sought to specifically define the role of cortical dynein activity in spindle bipolarity. Previous work has shown that the position, orientation, and oscillatory movement of the bipolar spindle within the cell is regulated, in part, by cortical dynein (59–61). Results from our model further indicate that cortical dynein activity impacts bipolar spindle length and promotes fluctuations in pole-to-pole distance over time (Fig. 5, 7). We used experimental techniques to validate this novel prediction made by our model. Fixed-cell imaging and analysis confirms that cortical dynein localization and activity impacts metaphase spindle length while live-cell imaging captures dynein-dependent fluctuations in bipolar spindle length over time (Fig. 8). We find that dynein-dependent changes in spindle length and dynamics are more robust in the absence of Nuf2 (in comparing Fig. 8 to Fig. S4 in the Supporting Material), suggesting that stable end-on kinetochore attachments may dampen dynamic changes in bipolar spindle length during mitosis. Future work could explore how this relationship is impacted in cells altered MT-kinetochore attachments stability, and define potential consequences of spindle length fluctuations, or lack thereof.

While mathematical models have extensively examined MT attachments to chromosomes and how these attachments drive chromosome movement and alignment during mitosis (16, 110, 111), we omit chromosomes and chromosome-derived forces in our model as end-on MT attachments to kinetochores are dispensable for bipolar spindle formation (33, 34). However, our data suggest that kinetochore-microtubule interactions may reinforce spindle stability. We find that the impact of cortical dynein disruption by Afadin depletion is more robust in cells lacking stable MT attachments to kinetochores (siNuf2) than in cells with robust MT attachments (siScr) (Fig. 8 B,D,E, Fig. S4 B,D,E). Furthermore, we find that short term dynarrestin treatment, which has been shown to partially disrupt kinetochore attachments (89), has a similar impact on cells regardless of Nuf2 activity (Fig. 8 B,D,E, Fig. S4 B,D,E). Together, these results suggest that MT attachments to kinetochores may impact spindle dynamics. Extension of our model to include chromosomes or chromosome-like structures and their associated forces would be necessary to define and test the contribution of chromosome or kinetochore-derived forces on spindle maintenance.

Failure in cell division or defects in chromosome segregation generates cells with abnormal, and sometimes double, the number of chromosomes and centrosomes. These defects contribute to tumor heterogeneity and cancer progression (112). Additional chromosomes pose a challenge for mitotic cells, as they may require larger spindles and are demonstrated to be sensitive to perturbation of mitotic motor proteins (113–119). We propose that dynamic changes in bipolar spindle length, driven by cortical dynein and/or activity, contributes to the spindle length requirements for chromosome capture and alignment, with particular relevance to cancer contexts where chromosome number is increased. Indeed, we see a significant increase in prometaphase cells following Afadin knockdown in RPE cells with stable kinetochore attachments (data not shown), suggesting defects in chromosome alignment when cortical dynein activity is lost. Future work can explore potential consequences of this phenotype.

While cortical dynein is dispensable for spindle bipolarity when Eg5 or HSET activity is unperturbed, our data implicate cortical dynein in achieving a maximum bipolar spindle length (Fig. 9 A). Our model predicts that high Eg5 activity can recover spindle length fluctuations with loss of cortical dynein, suggesting that this feature of spindle dynamics is not driven by cortical dynein specifically but instead reflects the stochastic balance between spindle forces. Further experimentation will be needed to confirm this prediction and to delineate the individual contributions of cortical dynein and Eg5 in this aspect of spindle bipolarity. Additionally, our model predicts that high cortical dynein activity is required for bipolar spindle formation when HSET activity is also high (Fig. 5 A(ii),B(ii),D, Fig. 10) (90, 91, 94). This may be particularly relevant in cancers cells with supernumerary centrosomes, where high HSET activity contributes to centrosome clustering, promoting bipolar spindle formation and continued cell proliferation (78, 90–96). High HSET levels have also been reported in cancer cells independent of centrosome number, although the functional implications of high HSET activity in this context remains unclear (90, 91). Our results indicate that when HSET activity is high, cortical dynein activity is required for bipolar spindle formation even when only two centrosomes are present. Therefore, we speculate that cancer cells having high levels of HSET, regardless of centrosome number, may be dependent on cortical dynein for bipolar spindle formation and accurate cell division. If true, therapeutic approaches to inhibit cortical dynein may be particularly effective at limiting mitotic progression in contexts of high HSET activity, independent of centrosome and/or chromosome number.

## AUTHOR CONTRIBUTIONS

Mercadante performed experiments and carried out all model simulations. Manning designed the experiments with Mercadante and Olson developed the code for simulations with Mercadante. All authors contributed to the writing of the article and data analysis.

## ACKNOWLEDGMENTS

Results in this paper were obtained in part using a high-performance computing system acquired through NSF MRI grant DMS-1227943 to WPI. We thank Neil Ganem for supplying the RPE GFP-Centrin cell line and the members of the Manning lab for feedback throughout the writing process. ALM is supported by a Smith Family Award for Excellence in Biomedical Research and DLM is supported by a NSF-GRFP. ALM, SDO, and DLM are further supported by NIH R01 GM140465-01.

## SUPPORTING MATERIAL

An online supplement to this article can be found by visiting BJ Online at http://www.biophysj.org.

## SUPPORTING CITATIONS

References (119-124) appear in the Supporting Material.

## Supporting Material

### ADDITIONAL MODEL DETAILS

#### Model Initialization and Algorithm

At the start of the simulation, random initial locations were chosen for centrosomes that matched the distribution from experimental data (Fig. 4 C). 300 MTs, randomly distributed between the two centrosomes, were initialized with angles and lengths *ℓ* chosen from a uniform distribution, *α* ∈ 𝒰[0, 2*π*) and *ℓ* ∈ [0, 0.5] and new MTs are nucleated at every time step (Fig. 2 A, Fig. S1 C,D). The states of each MT are updated based on the stochastic rules (e.g. Monte Carlo binding to cortical dynein if close to cortex based on binding probability 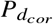). The states and configuration of the MTs then contribute to forces on each centrosome. The force-balance equation for each centrosome is solved to determine its new location. The MTs are then updated based on state (growing, shrinking, angle of vector direction changes). This is repeated at each time step until *t* = 30 min is reached.

#### Microtubules

MTs are nucleated at a rate *MT*_*nuc*_, undergo rescue (switch from shrinking to growing) at a rate *k*_1_ and undergo catastrophe (switch from growing to shrinking) at a MT-length dependent rate *k*_2_, defined as 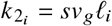, where *s* is a scaling factor, and *ℓ* _*i*_ is the length of the *i*^*th*^ MT (52). Sensitivity of the model (defined by spindle length) to the parameter *s* is shown in Table S4. While it has been well established that catastrophe frequency is MT-age dependent rather than length dependent (126), results from our model indicate that length and age are strongly correlated (Fig. S1 B in the Supporting Material). Following a standard Monte Carlo method, we choose *n*_2_ ∈ 𝒰 [0, 1] and the *i*^*th*^ growing MT undergoes catastrophe if 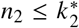, where 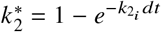 (13). Similarly, shrinking MT *i* will be rescued if 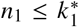 where 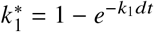. MTs that fail to undergo rescue depolymerize completely and are no longer considered in the system when *ℓ* _*i*_ ≤*v*_*g*_*dt*. The vector 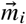 defines the direction of each MT *i*. While we do not account for physical bending of dynamic MTs, when defining the model terms and MT force generation by motor proteins, we account for the tendency of MT to bend, particularly at interpolar regions and scale the force as needed (Eqs. 6, 7, Fig. 2 B).

#### Model Analysis

While many parameters in our model have been well established by biological, biophysical, or mathematical studies, we define novel parameters in our model that we have optimized to reflect accurate spindle formation and maintenance. We explore the sensitivity of our model, as a readout of bipolar spindle length at t=10 min, with values above and below our selected parameters. While manipulating a parameter, all other parameters remain unchanged from the base case. Results are summarized in Tables S3 and S4.

To understand how model outcomes such as spindle length vary due to model stochasticity by MT dynamics and MT-motor protein binding and unbinding, we performed an increasing number of simulations with the same initial centrosome positioning. Traces of centrosome movement over time show different trajectories (results not shown), but the distance between centrosomes at t=25 min (spindle length) have similar trends, as shown in Table S2. All averaged simulation results reported are of a minimum of 10 simulations.

Simulations achieving spindle bipolarity were those having a spindle length of at least 17 *µ*m, as this is equal to our experimentally measured average bipolar spindle length in cells lacking stable chromosome attachments at t=25 (Fig. 3 C). Monopolar spindles were characterized by spindle length being less than half the average bipolar spindle length (8.5 *µ*m) at t=25 min.

#### Drag

While we maintain a constant drag coefficient in our model, we note that the drag on an object varies with size and proximity to the boundary (49, 58). We account for dynamic changes in drag by scaling motor-derived forces exponentially, with a strong dependence on the proximity of the centrosome to the point where the force is applied. Previously published results explored drag on a spherical object as it approached a boundary (58) and drag on an aster centered in the confined domain with a symmetric distribution of MTs of varying volume fraction (49). Inspired by these studies, we first defined a dynamic drag term as

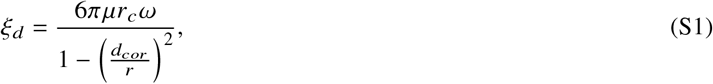

where *µ* is the viscosity of the cytoplasm (Table S1), *r*_*c*_ is the effective radius of the MT aster, calculated as the average length of MTs nucleated from centrosome *c, ω* is the volume fraction of MTs nucleated from centrosome *c, d*_*cor*_ is the minimal distance from the centrosome center to the cell cortex, and *r* is the radius of the cell (Table S1). We show that this drag coefficient increases as MT length and MT aster volume (MT density) increase (Fig. S2 A). Furthermore, *ξ*_*d*_ increases as the centrosome distance to the cell cortex decreases (Fig. S2 B). However, the drag term defined in Eq. (S1) did not capture the asymmetry of MT lengths and volume fraction throughout the 30 min of mitotic progression that we were modeling. Since MTs and forces are dynamic in our model, rather than applying a uniform drag coefficient on the centrosome, we define an exponential scaling term that is specific to each MT-motor interaction. We define this term as:

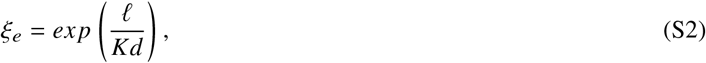

which considers *ℓ*, the distance from the centrosome center to the point where the force is applied, and *d*, the distance between the centrosome center and the object it is interacting with (either the cell boundary or the opposing centrosome). These terms account for the dynamic changes in drag described previously. To observe how this term impacts how force is felt by the centrosome center, we have a scatter plot of drag *ξ*_*e*_ on the *x*-axis, as a function of *ℓ*= *L*_*i*_ on the *y*-axis, the distance from the centrosome to the point where Eg5 and/or HSET bind. This plot is over a time course of 5 minutes in the simulation and has *d*_*cent*_, the distance between centrosomes, ranging from 4-15 *µ*m. We see that forces generated when *L*_*i*_ is large are correspond to a small *ξ*_*e*_, while when *L*_*i*_ is small, i.e. when the centrosome is close to where the force is being applied, *ξ*_*e*_ approaches 1 (Fig. S2 C).

**Table S1:** Parameter Values

| Parameter | Value | Description | Reference |
| --- | --- | --- | --- |
| **Microtubules** |  |  |  |
| $v_g$ | $0.183 \mu\text{ms}^{-1}$ | MT growth velocity (+ ends) | (62, 120) |
| $v_s$ | $0.3 \mu\text{ms}^{-1}$ | MT shrinking velocity (+ ends) | (62) |
| $v_b$ | $0.057 \mu\text{ms}^{-1}$ | MT shrinking velocity (+ ends)<br>bound to cortical dynein | (47) |
| $k_1$ | $0.167 \text{s}^{-1}$ | Rescue frequency | (62, 121) |
| $\kappa$ | $10 \text{pN}\mu\text{m}^2$ | Bending rigidity | Approximated (27, 47, 122) |
| $f_{\text{stall}}$ | $5 \text{pN}$ | Stall force of MTs | (123) |
| $MT_{\text{nuc}}$ | $2 \text{s}^{-1}$ | MT nucleation rate<br>per centrosome | (124) |
| $\theta$ | $10\pi/180$ | Slipping MT angle change | |
| **Motor Proteins** |  |  |  |
| *Dynein* |  |  |  |
| $f_{0,d}$ | $3.6 \text{pN}$ | Stall force of dynein | (37) |
| $v_{0,d}$ | $0.86 \mu\text{ms}^{-1}$ | Walking velocity of dynein | (37, 41) |
| $P_{d_{\text{cor}}}$ | $0.5$ | Probability of binding to<br>cortical dynein | |
| $P_{d_{\text{sp}}}$ | $0.1$ | Probability of binding to<br>spindle pole dynein | |
| $\mathcal{D}_d$ | $4v_g(dt) \mu\text{m}$ | Distance required for binding<br>to dynein | |
| $\mathcal{D}_{d_{\text{sp}}}$ | $1 \mu\text{m}$ | Distance required for binding<br>to dynein at spindle poles | |
| *Kinesin-5 (Eg5)* |  |  |  |
| $f_{0,Eg5}$ | $1.5 \text{pN}$ | Stall force of Eg5 | (40) |
| $v_{0,Eg5}$ | $0.2 \mu\text{ms}^{-1}$ | Walking velocity of Eg5 | (125) |
| $P_E$ | $0.5$ | Probability of binding to Eg5 | |
| *Kinesin-14 (HSET)* |  |  |  |
| $f_{0,HSET}$ | $1.1 \text{pN}$ | Stall force of HSET | (39) |
| $v_{0,HSET}$ | $0.2 \mu\text{ms}^{-1}$ | Walking velocity of HSET | (125) |
| $P_H$ | $0.5$ | Probability of binding to HSET | |
| $\mathcal{D}_{Eg5,HSET}$ | $v_g dt \mu\text{m}$ | Distance required for binding<br>to Eg5 or HSET | |
| **Other** |  |  |  |
| $r$ | $15 \mu\text{m}$ | Radius of the cell | |
| $c_r$ | $0.3 \mu\text{m}$ | Radius of a centrosome | |
| $\mathcal{D}_r$ | $2 \mu\text{m}$ | Distance for repulsive forces | |
| $K$ | $0.25$ | MT length-dependent scaling factor | |
| $C$ | $0.1$ | Antiparallel crosslinking scaling factor | |
| $s$ | $0.15 \mu\text{m}^{-1}$ | Scaling for catastrophe frequency | |
| $R$ | $1 \mu\text{m}$ | Scaling for repulsive forces | |
| $\mu$ | $0.7 \text{pNs}\mu\text{m}^{-2}$ | Viscosity of the cytoplasm | (56) |
| $\xi$ | $20.6 \text{pNs}$ | Drag coefficient | |
| $v_f$ | $0.0083 \mu\text{ms}^{-1}$ | Poleward flux | (68) |

**Table S2:** Average (Avg) Spindle Length for Simulations (Sims) at t=25 min.

| # of Sims | Avg length ($\mu\text{m}$) |
| --- | --- |
| 5 | 17.79 |
| 10 | 17.53 |
| 15 | 17.56 |
| 20 | 17.65 |

## SUPPORTING FIGURES AND MOVIES

**Movie M1** Simulation of the base condition, corresponding to Fig. 6B.

**Movie M2** Simulation with no MTs binding to cortical dynein 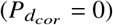, corresponding to Figs. 5 A(iii) and 7 B(i).

**Movie M3** Live-cell imaging of an RPE cell expressing GFP-centrin in a cell lacking stable MT attachments to kinetochores (siNuf2). Movie captured in a single z-plane at 60x at 15 sec intervals for 10 min, corresponding to Fig. 8.

**Movie M4** Simulation with no cortical dynein 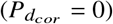 and high Eg5 (*P*_*E*_ = 1), corresponding to Fig. 9 D(iii).

**Figure S1:**
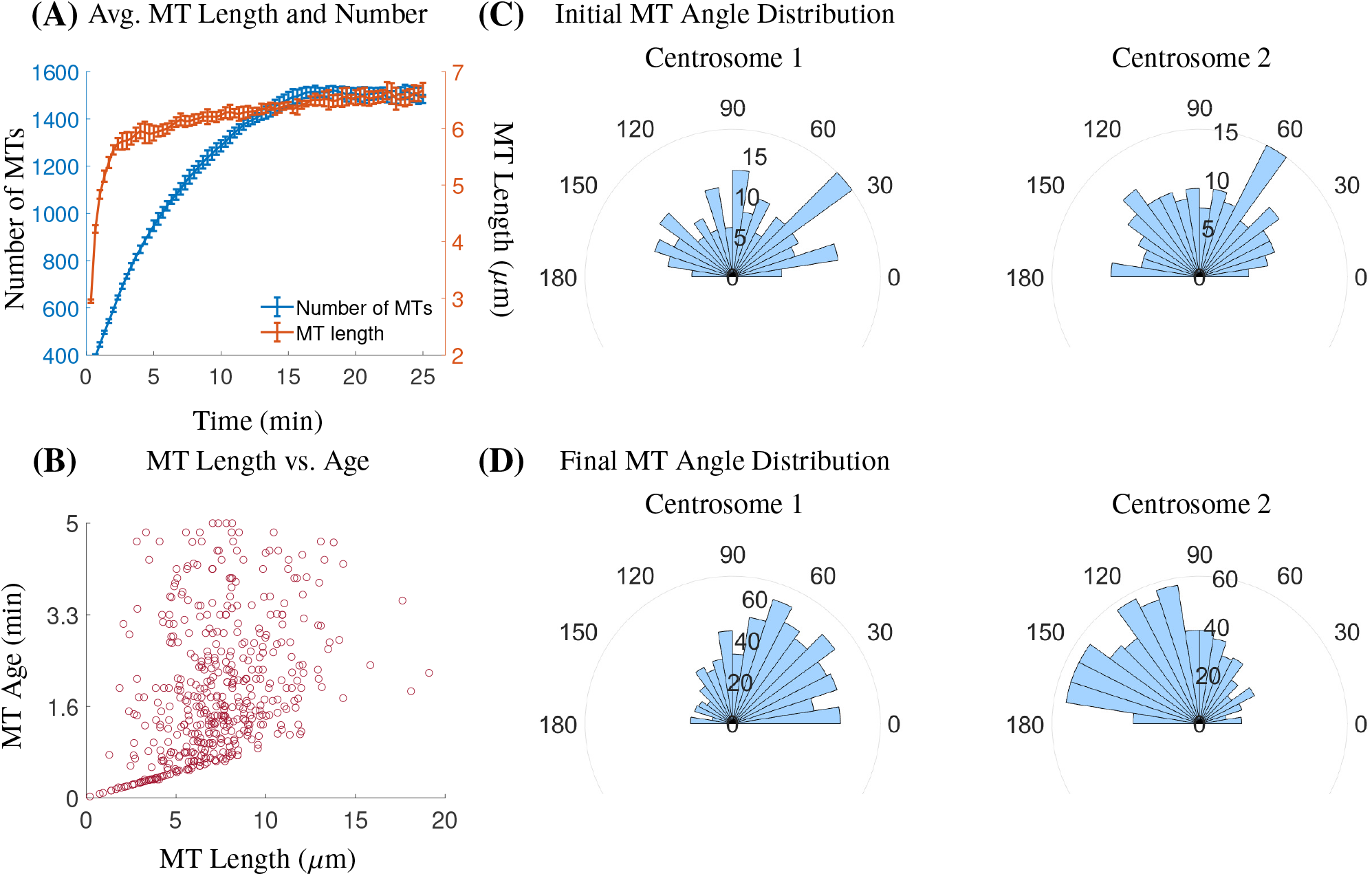
MT number, length, and distribution change throughout mitotic progression. (A) Average number of MTs and average MT length from 10 simulations of the base case of the model. Error bars are SD. (B) Scatter plot of MT length and age. (C) Representative histograms depicting the random initial MT angle distribution (in degrees) with respect to 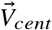 (t=20 sec) on both centrosomes from a single simulation. (D) Representative histograms depicting the asymmetric final MT angle distribution (in degrees) with respect to 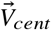 (t=25 min), with more MTs in the direction of 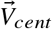 on each centrosome. Results from the same simulation are shown in (C) and (D).

**Table S3:** Sensitivity Analysis to crosslinking parameter *C* and force/drag scaling parameter *K*. Base case corresponds to *C* = 0.1 and *K* = 0.25. Results are averaged over 10 simulations at t=10 min.

|  |  |  |  |  |  |
| --- | --- | --- | --- | --- | --- |
| $C$ | 0.025 | 0.05 | **0.1** | 0.15 | 0.2 |
| Spindle Length ($\mu\text{m}$) | 5.09 | 14.8 | **16.5** | 17.31 | 17.67 |
| $K$ | 0.1 | 0.2 | **0.25** | 0.3 | 0.4 |
| Spindle Length ($\mu\text{m}$) | 11.43 | 14.88 | **16.5** | 18.42 | 20.35 |

**Table S4:** Sensitivity Analysis to MT catastrophe rate scaling *s* and constant drag parameter *ξ*. Base case corresponds to *s* = 0.15 and *ξ* = 20.6. Results are averaged over 10 simulations at t=10 min.

|  |  |  |  |
| --- | --- | --- | --- |
| $s$ | 0.05 | **0.15** | 0.3 |
| Spindle Length ($\mu\text{m}$) | 18.93 | **16.5** | 12.9 |
| $\xi$ | | **20.6** | 103 |
| Spindle Length ($\mu\text{m}$) | | **16.5** | 13.33 |

**Figure S2:**
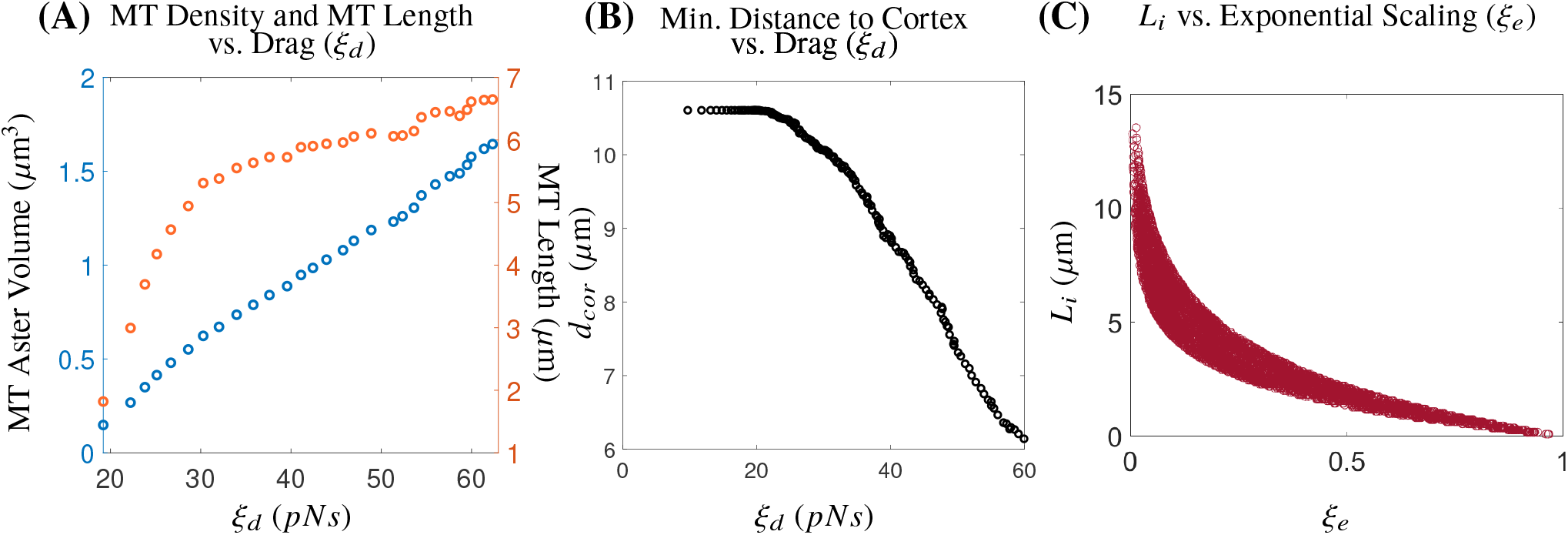
Exponential length scaling on forces captures increased drag due to MT length and MT density. (A) Scatter plot showing how increasing MT aster volume and MT length results in an increased drag coefficient (*ξ*_*d*_, shown on *x*-axis, Eq. S1) for a single simulation up to t=5 min. (B) Scatter plot of highlighting different centrosome distance to the cell cortex on the *y*-axis and dynamic drag coefficient (*ξ*_*d*_, Eq. S1) on *x*-axis. (C) Scatter plot of the exponential length scaling of motor-derived interpolar forces *ξ*_*e*_ (Eq. S2) on *x*-axis and the distance from the centrosome to the point of force application for all MTs bound to Eg5 and/or HSET (*L*_*i*_) on the *y*-axis.

**Figure S3:**
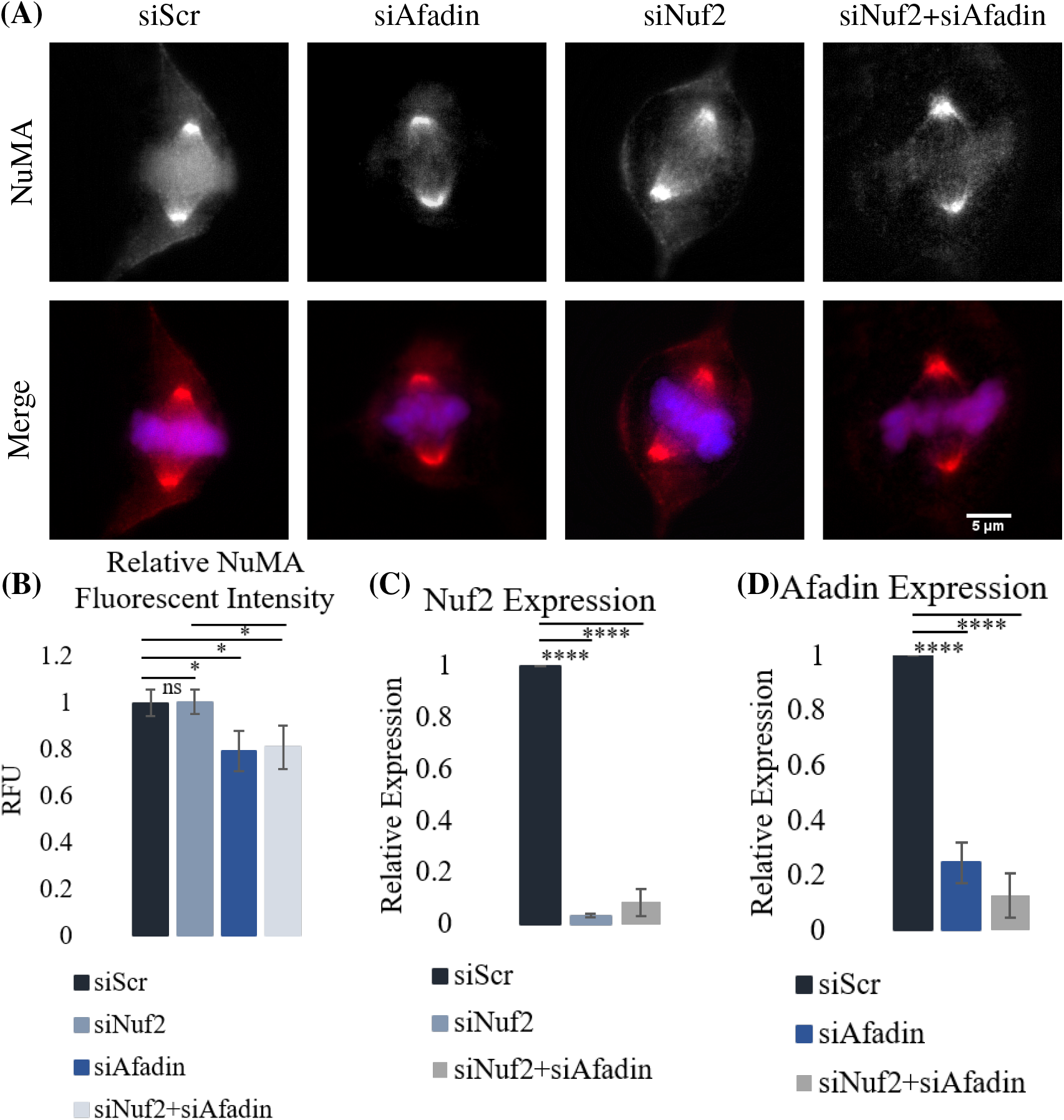
Afadin knockdown disrupts cortical NuMA localization during mitosis. (A) Immunofluorescent imaging of RPE cells following knockdown of Nuf2 and/or Afadin by siRNA. (B) Relative fluorescent intensity (RFU) of cortical-to-cytoplasmic NuMA. At least 20 cells were quantified for each condition from 3 independent replicates. (C) Quantification of Nuf2 RNA expression by qPCR. (D) Quantification of Afadin RNA expression by qPCR. Each condition was normalized to a control (siScr) and data is averaged over 3 independent replicates. Error bars are standard deviation. Significance was determined by student’s t-test (\**<*0.05, \*\*\*\**<*0.001, ns indicates not significant).

**Figure S4:**
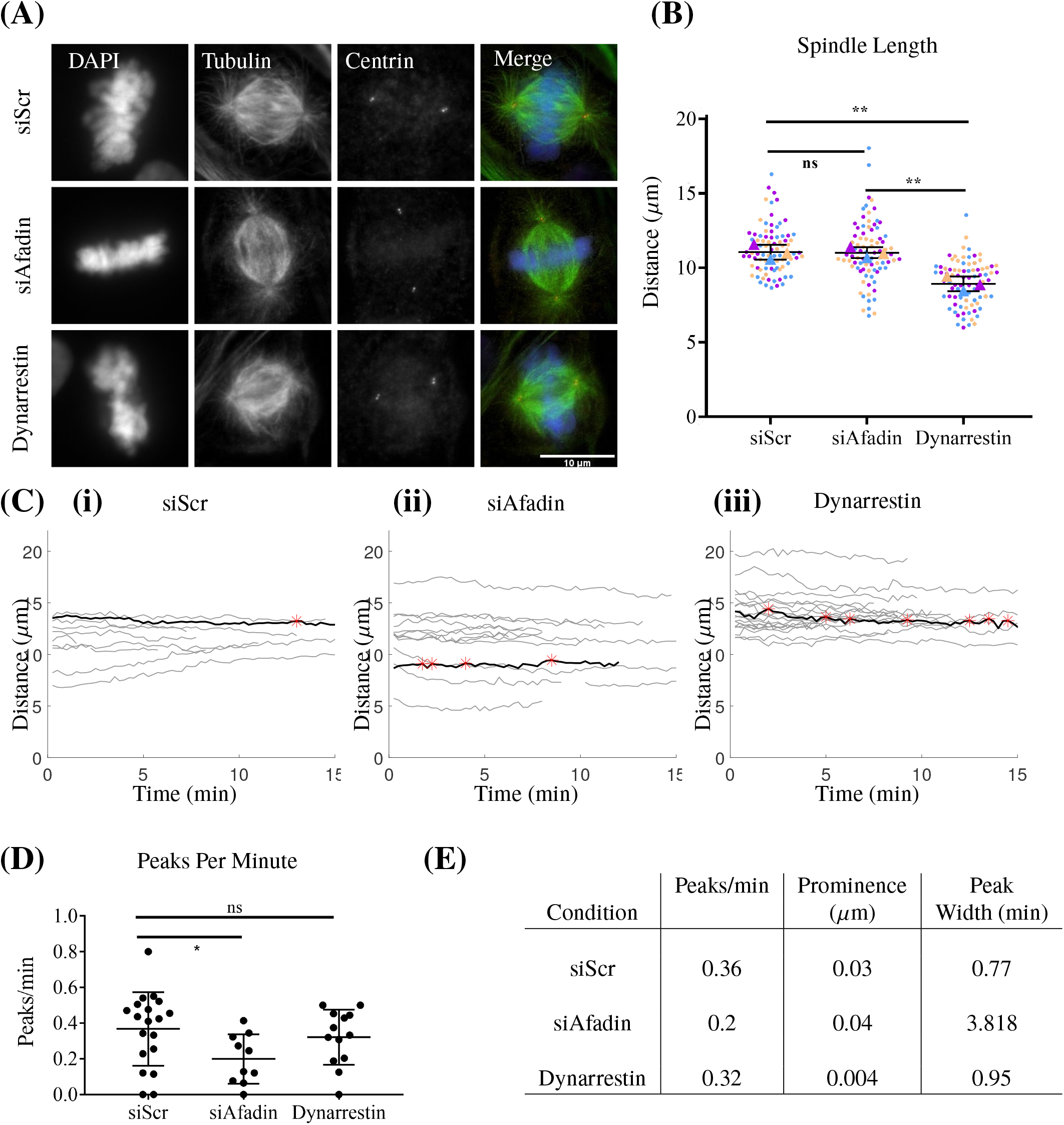
Fixed and live-cell imaging captures dynein-dependent changes in bipolar spindle length and spindle dynamics. (A) Fixed cell imaging of RPE cells stained for DAPI (DNA), Tubulin (MTs), and Centrin (centrosomes) in siScr, siAfadin, and Dynarrestin conditions. (B) Quantification of bipolar spindle length in siScr, siAfadin, and Dynarrestin conditions. Quantification performed on at least 25 cells from each condition for 3 biological replicates. Each color indicates a replicate and the average for each replicate is represented by a triangle of the same color. (C) Traces of spindle length over time of individual RPE cells expressing a GFP-centrin tag for siScr (i), siAfadin (ii), and Dynarresetin (iii) conditions. Red asterisks represent significant peaks for the curve shown in black. (D) Quantification of the average number of peaks per minute in siScr, siAfadin, and Dynarrestin conditions. Significance determined by one-way ANOVA. (E) Table showing the average number of peaks per minute, the average peak prominence, and average peak width from each condition. At least 10 cells were captured and quantified for each condition. All error bars are SD. *p*<*0.05 indicates statistical significance, ns indicates not significant.

